# Complete representation of action space and value in all striatal pathways

**DOI:** 10.1101/2020.03.29.983825

**Authors:** Moritz Weglage, Emil Wärnberg, Iakovos Lazaridis, Ourania Tzortzi, Konstantinos Meletis

## Abstract

The dorsal striatum plays a central role in motor and decision programs, such as the selection and execution of particular actions and the evaluation of their outcomes. A standard model has emerged where distinct output pathways encode separate motor-action signals, including selection-evaluation division in the matrix versus patch compartments. We used large-scale cell-type specific calcium imaging during motor and decision behaviors to determine and contrast the activity of individual striatal projection neurons (SPNs) belonging to one of the three major output pathways in the dorsomedial striatum – patch Oprm1+ SPNs versus the D1+ direct and A2A+ indirect pathway. We found that Oprm1+ SPNs were tuned to a number of different behavioral categories, such as to different movements, or to discrete actions and decisions in a two-choice task, and these complex representations were found to the same extent in all three striatal output pathways. The sharp tuning of individual SPNs was highly stereotyped over time while performing a specific task, but the tuning profile remapped between different behavioral contexts. In addition to action representations, SPNs showed pathway-independent representation of decision-variables such as the trial strategy and the action value. We propose that all three major output pathways in the dorsomedial striatum share a similarly complete representation of the entire action space, including task- and phase-specific signals of action value and choice.

## INTRODUCTION

The selection of specific actions is based on computations that produce prediction and evaluation of the action outcome, and proper action selection in each situation, is essential for the survival of all species. Action selection computations have been associated with neuron activity in basal ganglia circuits, where the striatum plays a central role in integrating information from cortical and subcortical circuits, which is then propagated to downstream targets for action execution (Cox and Witten, 2019; Gurney et al., 2001; Hikosaka et al., 2014). The striatum has been neuroanatomically divided into two major output pathways: the direct pathway targeting the globus pallidus interna (GPi) and the substantia nigra (SN), and the indirect pathway targeting the globus pallidus externa (GPe) (Kreitzer, 2009; Tepper et al., 2007). A circuit model has emerged based on the dichotomous organization of the striatum, where the direct and indirect pathway differentially control motor programs and explain the pathophysiology of movement disorders (Albin et al., 1989; Alexander and Crutcher, 1990; Nelson and Kreitzer, 2014). In this model, the striatal pathways regulate motor behaviors through antagonistic signals. The striatum has been shown to also encode key variables in decision-making such as action value (Lau and Glimcher, 2008; Samejima et al., 2005; Wang et al., 2013). In addition to the direct and indirect pathway, the striatum can be divided into compartments using neurochemical definitions (Graybiel and Ragsdale, 1978; Olson et al., 1972), notably the patch (also known as striosome) and matrix compartments, where the striatal patches exhibit high levels of mu opioid receptor (MOR) expression (Märtin et al., 2019; Pert et al., 1976). The striatal patches form a distinct pathway that projects to the GPi and SN (Fujiyama et al., 2011; Jiménez-Castellanos and Graybiel, 1989), and the function of patches has been primarily linked to the evaluation of actions (Friedman et al., 2015; White and Hiroi, 1998).

Distinct gene expression patterns have been used to genetically target, visualize, and manipulate the direct, indirect, and patch pathway (Gerfen and Surmeier, 2011; Gerfen et al., 1990; Gong et al., 2003). In support of a circuit model emphasizing the opposing function of the direct and indirect pathways, optogenetic manipulation has shown their differential role in reinforcement as well as action (Geddes et al., 2018; Kravitz et al., 2012; Tai et al., 2012). In contrast, concomitant activation of both direct and indirect pathways during movements suggests a possibly mixed motor representation (Cui et al., 2013), and the direct and indirect pathways show complementary roles in action sequences (Tecuapetla et al., 2016). Imaging of neuron activity in the direct and indirect pathway (Klaus et al., 2017), in striatal patches (Bloem et al., 2017; Yoshizawa et al., 2018), and in mouse models of movement disorders (Parker et al., 2018), has together provided some supporting evidence for the opposing role of striatal pathways for simple motor and action behaviors, but has also challenged the unitary representation of behavioral events.

To address the role of the striatal patches as well as the other two major output pathways of the dorsomedial striatum in the representation of motion, actions, and decision-making variables, we recorded the activity of single SPNs belonging to the patch (Oprm1+) pathway as well as the direct (D1+) and indirect (A2A+) pathway using transgenic mice performing specific behaviors. Our findings on the activity of Oprm1+ SPNs during locomotion versus decision-making reveal a context or task-specific encoding of the discrete movements during exploration, the discrete actions and task strategy (action space and value) in a two-choice task, that form a complete representation of the action space and value that is shared with the major direct and indirect striatal output pathways.

## RESULTS

### The activity of dorsomedial D1+, A2A+, and Oprm1+ SPNs during locomotion

To define the role of the patch Oprm1+ SPNs during locomotion versus action selection and evaluation and to compare with the two other major output pathways of the dorsal striatum, we imaged the genetically-encoded calcium sensor GCaMP6s in SPNs through implanted GRIN lenses targeting the anterior part of the dorsomedial striatum in the right hemisphere of freely moving mice (Fig. 1A and Fig. S1). To map the activity of Oprm1+ SPNs, we used an Oprm1-Cre mouse line that specifically labels Oprm1+ cells (Märtin et al., 2019). We imaged the calcium signal in individual SPNs belonging to the three major output pathways, defined genetically by the expression of D1 (i.e. D1-Cre for direct pathway), A2A (i.e. A2A-Cre for indirect pathway), or Oprm1 (i.e. Oprm1-Cre for patches) (Fig. 1B). We applied the CNMF-E algorithm to extract regions-of-interest (ROIs) and deconvolved the calcium traces for single neurons (Giovannucci et al., 2019) to better capture the physiological range dynamics of the neuron activity (Fig. 1C).

**FIGURE 1.**
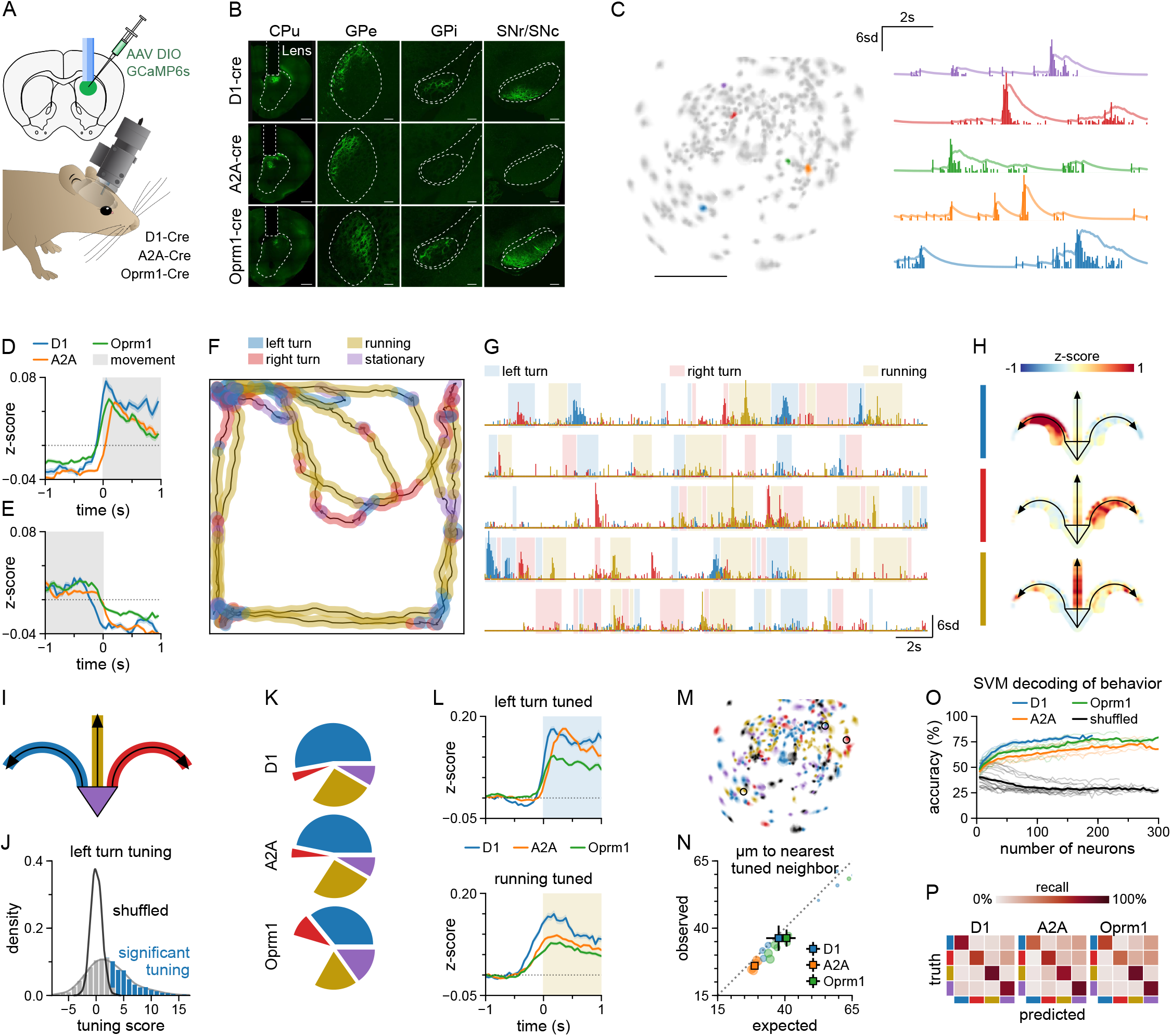
The activity of dorsomedial D1+, A2A+, and Oprm1+ SPNs during locomotion. A) Illustration of experimental approach to image neuron activity in SPNs of the dorsomedial striatum (DMS). A Cre-dependent AAV vector was injected into D1-Cre (direct pathway), A2A-Cre (indirect pathway), or Oprm1-Cre (patch pathway) mice (top). Mice were implanted with a GRIN lens. Calcium signals were recorded with head-mounted miniscopes (bottom). B) Images show Cre-dependent GCaMP6s expression (green) in caudate putamen (CPu), globus pallidus externa (GPe), globus pallidus interna (GPi), and substantia nigra (pars reticulata and compacta, SNr/SNc). Scale bars: 1 mm in the first column, 200μm in the other panels. C) Example of neurons detected in the field-of-view of a single recording session by the CNMF-E algorithm (left map). Calcium signals from the five neurons color-coded in the map. Transparent lines show denoised fluorescence and opaque bars the deconvolved signal. Scale bar: 200μm. D) Population average activity aligned to movement onset. Lines show mean, shaded areas ±SEM; n=632 D1-neurons from 7 sessions, 1604 A2A-neurons from 6 sessions and 1465 Oprm1-neurons from 10 sessions. E) Population average activity aligned to movement offset. Lines show mean, shaded areas ±SEM; n=632 D1-neurons from 7 sessions, 1604 A2A-neurons from 6 sessions and 1465 Oprm1-neurons from 10 sessions. F) Example showing the behaviors detected in the open field arena (2 minutes, Oprm1+ mouse). The behavior was segmented into left turns, right turns, running bouts and quiescence. Color-code same as in panel G. G) Deconvolved signals for three selected neurons in the patch pathway (Oprm1+ mouse, behavior shown in F). H) Heatmaps of the average responses of the three neurons in G during the four segmented behaviors. Arrows represent the three movements, the triangle quiescence. I) Open field task schematic indicating the color coding of the segmented behaviors. Same color code used in panels K, M and P. J) Histogram of the left turn tuning scores of all Oprm1+ neurons (n=1465 from 10 sessions). K) The proportion of neurons of each primary tuning for all three Cre lines. Color code follows I. n=632 D1-neurons from 7 sessions, 1604 A2A-neurons from 6 sessions and 1465 Oprm1-neurons from 10 sessions L) Average activity of left turn and running-tuned populations around the onset of the respective behavior. n(left turn) = 514 tuned D1-neurons from 7 sessions, 1324 tuned A2A-neurons from 6 sessions, 1210 tuned Oprm1-neurons from 10 sessions. n(running) = 330 tuned D1-neurons from 7 sessions, 1032 tuned A2A-neurons from 6 sessions, 737 tuned Oprm1-neurons from 10 sessions. M) Example field of view illustrating the spatial distribution of individual neurons color-coded according to their primary tuning. Oprm1+ mouse, color code follows I. N) The observed average distance of each neuron significantly tuned to a behavior to its closest neighbor tuned to the same in comparison to the average minimum distance expected by chance (computed based on repeated shuffles of neuron identity). Circles: single session averages, radii proportional to the number of recorded neurons. Squares: average value for each Cre-line, sessions weighted by the number of neurons. Error bars: ±SEM (bootstrapped). N=7 sessions from D1-Cre mice, 6 sessions from A2A-Cre mice, 10 sessions from Oprm1-Cre mice. O) Decoding accuracy of support vector machines (SVMs) trained to predict the current behavior from neuron activity. Thin lines: single sessions; thick lines: average for each Cre-line. Black lines: behavior labels segment-wise shuffled. N=7 sessions from D1-Cre mice, 6 sessions from A2A-Cre mice, 10 sessions from Oprm1-Cre mice. P) Average confusion matrix of SVM predictions. All of a session’s neurons were used to train the SVM; confusion matrices were averaged with the population size as weights. Behaviors are color coded following I. N=7 sessions from D1-Cre mice, 6 sessions from A2A-Cre mice, 10 sessions from Oprm1-Cre mice.

To first map the activity of individual SPNs during basic motor programs, we exposed mice to an open field arena and recorded the activity of SPNs during exploration and self-paced locomotion. We found on the population level that the Oprm1+ pathway as well as the D1+ and A2a+ pathway showed on average a bias for encoding contralateral turns, and an increase in activity upon movement initiation and a decrease after stopping (Fig. 1D-E and Fig. S2). To map in more detail the SPN activity during specific behavioral events, such as turning, acceleration, start-stop, we extracted a set of discrete behaviors in the open field arena (e.g. left turn, right turn), and plotted first the activity of selected Oprm1+ SPNs to map their tuned responses. We found many examples of Oprm1+ SPNs that showed representation of simple movement categories, such as left versus right turning or forward acceleration (Fig. 1F-H and Fig. S2). To directly compare the activity features for SPNs belonging to the three output pathways, we computed tuning scores for the main behavioral categories for all recorded SPNs. The distribution of tunings was unimodal for all behavioral categories, but wider than expected by chance (Fig. S2). Based on these distributions, we defined the significantly tuned neurons (Fig. 1I-J). Overall, we found significantly tuned neurons that represented for example left turns (contralateral to the recording side) or running speed. Importantly, we found a surprising similarity between the D1+, A2A+, as well as Oprm1+ pathways in term of the proportion of tuned neurons for each of the different behavioral categories as well as in the dynamics of the event-related calcium signal (Fig. 1K-L, Fig. S2). Similar to a previous study on the activity of D1+ and D2+ SPNs in dorsolateral striatum during locomotion(Klaus et al., 2017), we found that neurons of a specific tuning type were slightly closer in space than expected by chance for all three pathways but they did not form clear spatial clusters (Fig. 1M-N). To further assess to what extent the recorded neuron activity in each striatal pathway contained information to encode the different behavioral categories in the open field, we trained support-vector machines (SVMs) on the neuron activity data in order to decode the different behaviors. We found high decoding accuracy for the mouse behavior using neuron activity data from SPNs belonging to either the D1+, A2A+, or the Oprm1+ pathway, and decoding was not significantly different for data from the three pathways (Fig. 1O-P and Fig. S7B). This supports that Orpm1+ SPNs in dorsomedial striatum can encode the various parameters of the motor program, and that the three major striatal output pathways all contain a similarly complete representation of the different behavioral categories, formed by the wide range of tuning profiles of individual SPNs found in each of the direct, indirect, and patch pathway.

### The D1+, A2A+, and Oprm1+ SPNs represent the action space of a two-alternative choice task

Since striatal patches have been proposed to primarily guide reward-based behavior and the evaluation of actions, we investigated the tuning profile of SPNs in a behavioral context of decision-making and action value. We imaged SPN activity in a behavioral task that requires evaluation of actions and updating of predictions regarding action outcome, based on a modified two-alternative choice task (Tai et al., 2012). In this task, mice were trained to freely initiate a trial by nose-poking into a center port, which allowed them to then choose to nose-poke on the right or left side port to receive a liquid reward (Fig. 2A). The probability of reward delivery in the one of the side ports (i.e. the correct port) was set to 75%. After a reward was delivered, there was a 5% chance that the reward port switched side. Importantly, mice were not given any cue to indicate the rewarded port, forcing them to keep track of their actions and update their choices based on recent trial outcome history. We trained mice in the task for at least 3 weeks, resulting in correct port choice in 67% of the trials, indicating that the mice learned to dynamically adapt their choices to trial outcome (Fig. 2B-C). Mice showed appropriate switching behavior, by updating their choice based on the reward history, showing a win-stay (Fig. 2D) and lose-switch strategy (Fig. 2E).

**FIGURE 2.**
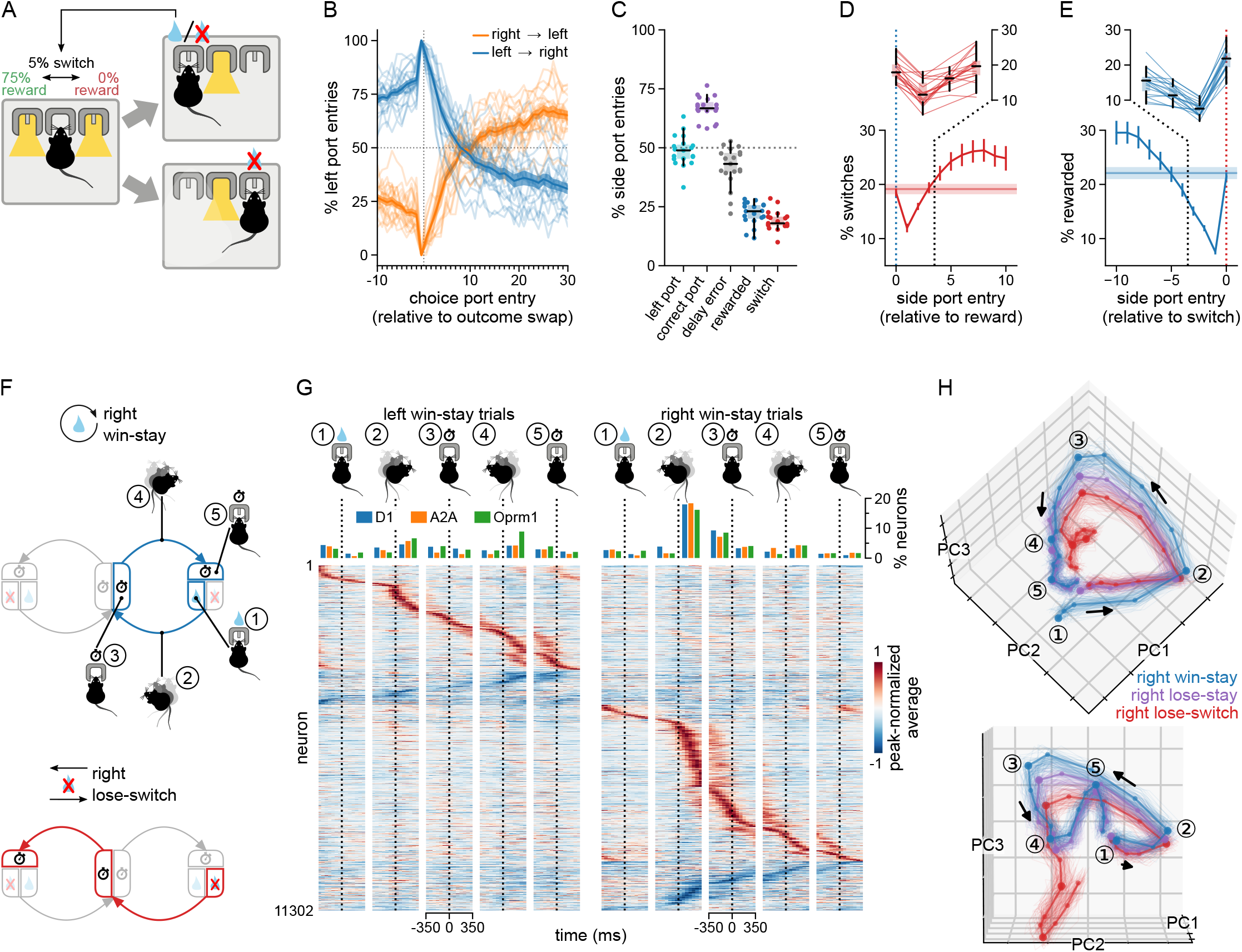
The D1+, A2A+, and Oprm1+ SPNs represent the action space of a choice task. A) Illustration of the 2-choice switching task. Mice initiated a trial by poking the central port, followed by a choice for one of the side ports. Only one side port delivered sucrose rewards (reward probability of 75%). 5% of reward deliveries were followed by a covert switch of reward delivery side. To initiate trials or receive rewards, mice had to stay a minimum of 350 ms in the respective ports. B) Mice successfully shifted their port preference after covert switches of the rewarded port. A switch of the reward from the right to the left port was followed by a gradual increase in the fraction of left port entries on subsequent trials (red line), a left to right switch vice versa (blue line). Reward port switches only occurred after rewarded choices. n = 3167 switches. Thin transparent lines: individual mice. C) Key statistics of the behavior. *Correct port*: entry into the port currently assigned the reward. *Delay error*: failure to trigger a choice port due to premature withdrawal (<350ms) from same or the initiation port. *Switch*: choice of the port not selected last. n=299083 trials from 271 sessions from 19 mice. Transparent dots: individual mice. D) Win-stay behavior: The probability of a choice switch was lowest immediately after a rewarded choice (side port entry 0). Horizontal line: overall average fraction of switch choice port entries. Thin transparent lines: individual mice. E) Lose-switch behavior: Mice were least likely to obtain rewards (i.e. were most likely to lose) on choice port entries preceding a switch choice (side port entry 0). Horizontal line: overall fraction of rewarded choice port entries. Thin transparent lines: individual mice. F) Illustration of “win-stay” (top) and “lose-switch” (bottom) trials after right side port outcomes. The outcome phase was treated as the start of the trial to reflect the dependence of a stay or switch choice on the outcome obtained last; a win-stay trial starts with a win (reward), a lose-switch trial with a loss (no reward). The task space was defined by twelve trial phases. Rectangles correspond to time spent in the center or outcome ports (left, right). Arrows represent movements between the ports. The center port is split into two phases according to the direction of the upcoming choice. Side ports are subdivided into the delay period (clock in upper rectangle), and reward (drop, inner part) and omission (x, outer part) phases. G) The average response of neurons to the phases of left and right port win-stay trials, centered on the phase onset times (time 0, window: ±350 ms). Neurons were peak-sorted. The bar charts indicate the percentage of neurons of each pathway (color-coded) with a positive peak falling within the 350 ms time window in the raster below. Pooled data from all recordings. H) Pseudo-trial activity trajectories based on single trial data pooled over all neurons recorded. The trajectories separate by trial type (color-coded), appearing ordered according to value implicit in the strategic choice made (win-stay > lose-stay > lose-switch). Trials were binned in scaled time (4 bins per phase). Trials were drawn in random order for every neuron. D1+ n=376; A2A+ n=1663; Oprm1+ n=1864 neurons; from 5, 5, 16 sessions, respectively. Mean trajectories of the pseudo-trial data used for dimensionality reduction (PCA; thick lines) and resampled pseudo-trials (thin lines). Error bars and shading: ±SEM. Boxplot whisker range: upper (lower) quartile to highest (lowest) value within 1.5 IQR

To define whether neuron activity reflected the strategy employed in the different trial types, we first focused our analysis on win-stay trials (e.g. reward in the right port followed by choice for the right port) versus lose-switch trials (e.g. no reward in the right port followed by choice for the left port) (Fig. 2F). We plotted the activity of individual SPNs from D1+, A2A+, and Oprm1+ pathways in the five discrete phases of the win-stay trial. We found that the peak activity of all recorded neurons across all win-stay trials tiled the entire task structure, suggesting that SPN activity did not show a clear bias or structure towards certain phases of the trial regardless of pathway identity (Fig. 2G).

To further explore the relation between neuron activity, trial structure and behavioral strategy, we pooled single-trial recording data of all trials following a right choice from several recording sessions to construct one pseudosession, which comprised approximately 20-30 pseudotrials of each trial type. We then analyzed the single-pseudotrial activity data by performing a dimensionality reduction to extract task-relevant principal components (principal component analysis, PCA), and we found that neuron activity contained structured information to describe the entire task trajectory, but also to discriminate the trial type (win-stay, lose-stay, lose-switch; Fig. 2H). The neuron activity therefore encodes two critical aspects for proper task performance: the sequential representation of phases in the trial as well as the decision variables differentiating the main trial type strategies.

### The D1+, A2A+, and Oprm1+ SPNs share a similar representation of the trial phases

The widely distributed activities of SPNs over many trial phases lead us to investigate the SPN activity during the different phases of the trial, and we focused on how individual neurons in the D1+, A2A+, and Oprm1+ pathways were modulated by discrete trial phases. We found that some Oprm1+ neurons were highly tuned to discrete phases in the task, representing specific movements (e.g. center to right turns) as well as outcome and choice signals (Fig. 3A-B). Furthermore, we found examples of Oprm1+ SPNs with sharp tuning for all the other trial phases (Fig. S3). To investigate and compare the tuning profile of all the striatal pathways, we calculated tuning scores for neurons in each output pathway for all twelve discrete task phases (Fig. 3C-D and Fig. S4). We found that D1+, A2A+, and Oprm1+ pathways showed a similar proportion of tuned neurons for each trial phase, and that all pathways showed a tuning bias for trial phases representing contralateral movements (Fig. 3E). We selected the significantly tuned neurons for the four different movements in the task and visualized their average activity during each phase. We found that the average amplitude was similar and phase-specific for all pathways (Fig. 3F). SPNs tuned to the reward phase were not modulated by reward magnitude (Fig. S5). Importantly, the tuning of single neurons was often not limited to a single trial phase, and the tuning profiles did not form well-separated clusters, suggesting a rich representation of the trial (Fig. 3G and Fig. S6). We calculated the spatial organization of neurons that share a significant tuning and found that these neurons were closer than expected by chance but did not form distinct clusters, further supporting the absence of clear functional segregation of SPNs in space (Fig. 3H-I).

**FIGURE 3.**
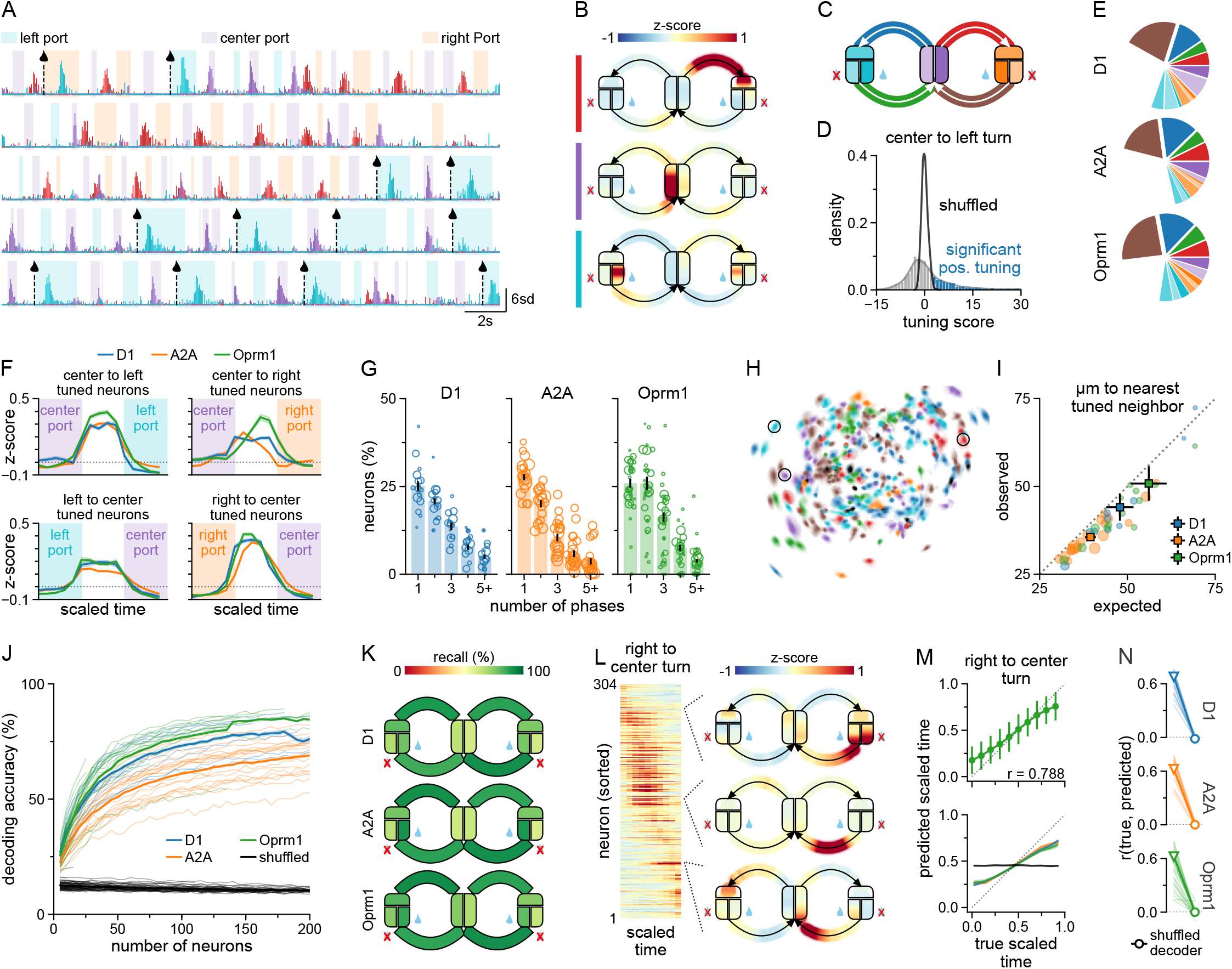
The D1+, A2A+, and Oprm1+ SPNs share a similar representation of the trial phases. A) Deconvolved traces from three example neurons, respectively tuned to the center-to-left turn (red), the center port delay phase which precedes a left turn (purple), and left port reward delivery. Shading indicates port occupancy. Black dashed lines and drop pictograms mark reward delivery. B) The average activity of three example neurons in every trial phase over the entire recording session plotted as heatmaps structured according to the 2-choice task schematic. The neurons are the same as in panel A; color bars indicate correspondence. Trial-level phase activity scaled to uniform length. C) Illustration of the 2-choice task schematic divided into twelve phases. Rectangles correspond to time spent in the center or outcome ports (left, right). Arrows represent movements between the ports. The center port is split into two phases according to the direction of the upcoming choice. Side ports are subdivided into the delay period (upper part), and reward (drop, lower inner part) and omission (x, lower outer part) phases. Reference for phase color-coding in panels E, F and H. D) Histogram of the center-to-left movement tuning scores of all Oprm1+ neurons (n=2793). The tuning scores were computed by z-scoring neurons’ mean activity with the mean and standard deviation of a sampling distribution obtained using block-wise shuffled behavior data. Significance: mean activity above the 99.5th percentile. E) Pie charts proportioned according to neurons’ primary phase-tuning, per Cre-line. Colors follow C. n=1943 D1-neurons, 6566 A2A-neurons, 2793 Oprm1-neurons. F) Average activity of 4 of the 12 populations defined by phase-tuning (the movements) during the phase they are tuned to. n(center-to-left neurons) = 413/1517/751; n(center-to-right neurons) = 279/928/426; n(left-to-center neurons)=228/883/424; n(right-to-center neurons)=581/1785/996 (D1/A2A/Oprm1). G) Bar plots of the weighted mean percentage of neurons significantly tuned to several counts of phases, by Cre-line. Sessions were weighted by the number of neurons. Circles: single sessions, radii proportional to number of neurons. Error bars: ±SEM (bootstrapped). H) Oprm1+ single-cell ROIs for an example session, color-coded by primary tuning. Black: no significant tuning. Circles mark ROIs whose activity is shown in A and B. I) Similarly tuned neurons are marginally closer in space than expected by chance (N=16 D1-sessions, 19 A2A-sessions, 30 Oprm1-sessions). J) Decoding accuracy (fraction correctly predicted phases) for randomly sampled SPN sub-populations of increasing size. Thin lines: single sessions. Thick lines: average per Cre-line. N=16 D1-sessions, 19 A2A-sessions, 30 Oprm1-sessions. K) The SVM decoding recall by phase when trained on the full population of each session. The average recall was weighted by the number of neurons in each session. L) Sequential neuron activity through a task phase. Raster plot of the average activity for all neurons recorded in an example session (Oprm1+, right port to center port movement). Neurons sorted by the peak of the activity (left panel). Heatmaps show three example neurons across the entire task space in the same session (right panel). M) The trial phase substructure can be decoded from neuron activity. Top panel: the actual versus the decoded time progression through the phase (right port to center port turn) for the session shown in L. Bottom panel: the actual versus decoded progress through the phase averaged over all sessions. Individual sessions weighted by the number of recorded neurons; Cre lines color coded as in N. Black line: decoder trained on shuffled data. Shaded area: ±SEM (bootstrapped). N) The correlation between true and predicted progress through right-to-center turns for all sessions (thin lines), compared to the correlation from shuffled data. Thick lines indicate averages, weighted by the number of recorded neurons in each session. Error bars: ±SEM (bootstrapped).

The rich tuning profile found in the D1+, A2A+, and Oprm1+ pathways suggested that neuron activity should include enough information to categorize the trial phase on a single trial basis. To test this, we trained SVMs to predict trial phases based exclusively on the neuron activity. We found that activity from even a relatively small fraction of the neurons was sufficient to decode any of the twelve task phases with high accuracy, and that the decoding accuracy was similar for neuron activity from the D1+, A2A+, and Oprm1+ pathways (Fig. 3J-K and Fig. S7). The high accuracy of these predictions demonstrates that all three pathways form trial or action space representations that are conserved on a trial-by-trial basis. Furthermore, we found clear examples where the peak neuron activity tiled the subparts of a single phase, exemplified by the peak activity of Oprm1+ SPNs during a right port to center port turn, which suggested that SPN activity contained information on the phase substructure in a trial (Fig. 3L). To determine whether neuron activity could represent the detailed temporal organization of phase progression, we trained SVMs on the activity of D1+, A2A+, and Oprm1+ pathways and found that we could accurately predict the temporal substructure of the turning behavior (right-center port turn) over the entire phase for all three pathways (Fig. 3M-N and Fig. S7).

### The phase tuning of D1+, A2A+, and Oprm1+ SPNs is task-specific and highly conserved across sessions and days

We next investigated whether the tuning profile of individual SPNs representing discrete behaviors in the open field was conserved in the choice task during similar actions (e.g. turning, stopping). We therefore defined whether SPNs showed comparable tuning during similar movements across the two different contexts and environments. To track individual SPNs across different behavioral contexts, we recorded the activity of single neurons from the D1+, A2A+, and Oprm1+ pathways first in the open field and immediately after in the two-choice task, without removing the miniscopes to ensure accurate tracking of neuron identity. We found that Oprm1+ SPNs with significant and highly stereotyped tuning for example to a left turn in the open field remapped their tuning profile in the choice task and instead represented the right-center port movement (Fig. 4A-C and S8). Similarly, a sharply left-center port tuned Orpm1+ SPN in the choice task showed no significant tuning in the open field (Fig. 4D-F and S8). When we compared the tuning of neurons for left or right turns between the open field and the two-choice task, we found that tuning preference between the two behavioral contexts were largely independent (Fig. 4G). Supporting the non-conserved tuning profile of SPNs between the two behavioral contexts, when comparing the primary tunings of all SPNs between the open field and the two-choice task we found that the tuning identity of SPNs was not conserved across the two contexts (Fig. 4H and Fig. S8). Since remapping of SPN tuning was evident between behavioral contexts within a single day, we further investigated whether the tuning profile within a single behavioral context was stable across sessions (i.e. over several days), we tracked the activity of single neurons in the choice task over days to weeks (Fig. 4I). We found that the significantly phase-tuned neurons maintained their tuning profile across several recording sessions, even for very sharp phase tunings (Fig. 4J and Fig. S8). Importantly, the conservation of SPN tunings across days was confirmed as we could use SVMs trained on neuron activity from one session to predict the trial phase in sessions from subsequent days (Fig. 4K). In summary, we found that the tuning profile of SPNs remapped in a context-specific fashion, thereby forming complex action representations that are stable and unique for each behavioral context.

**FIGURE 4.**
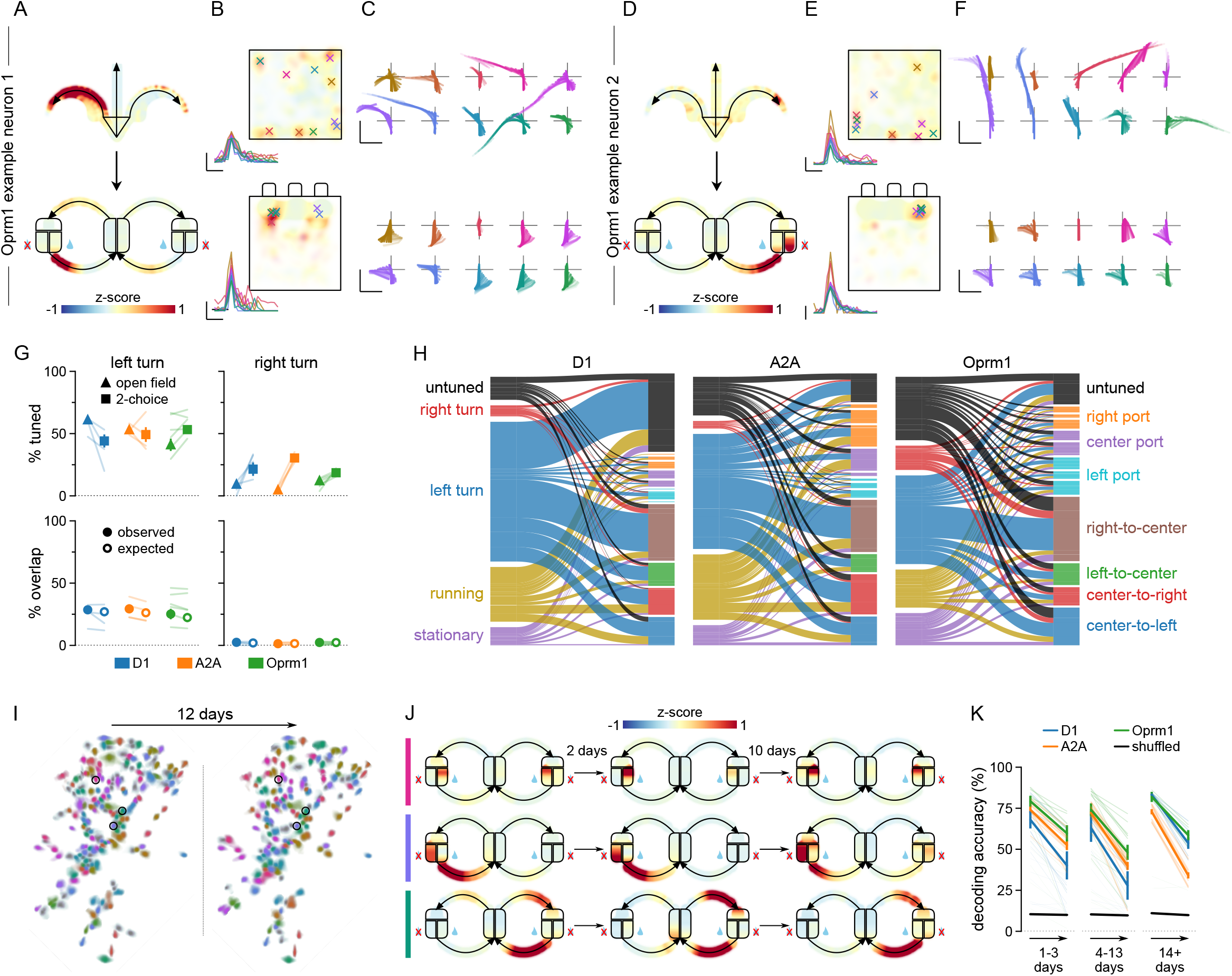
The phase tuning of D1+, A2A+, and Oprm1+ SPNs is task-specific and highly conserved across days. A) Example of Oprm1+ neuron with remapped tuning in the open field and the 2-choice task. The neuron is tuned to left turns in the open field and to right turns (left-center port). Imaging was performed in the two tasks without detaching the miniscope. B) Spatial heatmaps of the activity of the neuron shown in A in the open field arena (top) and the 2-choice operant chamber (bottom). Crosses mark the top 10 most prominent deconvolved calcium events in the respective recordings; events shown in the lower left corners of the maps. Scale bars: 10 std, 250 ms. C) Movement trajectories of the animal during the top 10 most prominent calcium events shown in B. Rotated such that the animal initially faces upwards. The animal’s posture is represented frame-for-frame by lines connecting the base of the tail, the center of the body, and the point between the ears. Scale bars: 5 cm, 5cm. D) Example of Oprm1+ neuron recorded in the same session as neuron in A-C. Neuron does not show tuning to any of the tracked movements in the open field, but responds sharply during left turns (right-center port) in the 2-choice task. E) Spatial heatmaps of the activity and the top 10 most prominent calcium events of the neuron shown in D in the open field arena (top) and the 2-choice operant chamber (bottom). F) Movement trajectories of the animal during the top 10 most prominent calcium events. G) Top left: Percentage of neurons significantly positively tuned to left turns in the open field and 2-choice tasks (tuned to either the center port to left or the right port to center turns), respectively. Bottom left: Comparison of the observed and randomly expected percentage of neurons tuned to left turns in both tasks. Panels in the right column show the same quantifications for right turn-tuned neurons. Sessions weighted by the number of recorded neurons (N=4 D1-sessions, 4 A2A-sessions, 6 Oprm1-sessions). Error bars: ±SEM (bootstrapped). H) Alluvial plot showing how the primary tunings in the open field (left side) change in the 2-choice task (right side). n=406 D1-neurons, 1111 A2A-neurons and 985 Oprm1-neurons were followed. I) Spatial filter map of an example recording session (Oprm1-Cre) and aligned filters from a session recorded 12 days later. The depicted filters were transformed by the registration process. Colors are chosen arbitrarily, but consistent for aligned neurons. Gray filters are without a match. J) Trial heatmap of 3 neurons matched across 3 sessions, including those shown in C. The colored bars match the color of the respective, circled filters in C. K) SVMs decode the trial phase across sessions (color code shows pathway). Each pair of sessions (training session + test session) is shown with a transparent line, indicating the cross-validated accuracy in the training session (left) and the test session (right). The thickness of transparent lines is proportional to the number of aligned neurons for that pair. Opaque lines correspond to the average decoding accuracy across pairs of sessions, weighted by the number of aligned neurons. SVM decoding accuracy on shuffled data (black). N(1-3 days) = 65/55/85 session pairs; N(4-13 days) = 25/60/85 session pairs; N(14+ days) = 25/30/25 session pairs (D1/A2A/Oprm1). Error bars: ±SEM (bootstrapped).

### D1+, A2A+, and Oprm1+ pathways represent the task strategy and the action value

The tuning of neurons in the D1+, A2A+, and Oprm1+ pathway to specific trial phases revealed a structured activity pattern that represented the entire trial structure. In addition to the representation of the trial phases, mice in the choice task also needed to keep track of more abstract task variables, such as the value and outcome of actions. To map the neuron activity that differentially represents the variables that match value and decision aspects of the task, we first identified neurons with activity patterns that showed modulation by the trial outcome and subsequent choice, which define the win-stay versus lose-switch strategy. We found examples of Oprm1+ SPNs that showed modulation by trial type, either increasing their activity during lose-switch decisions or increasing activity in win-stay strategies (Fig. 5A and Fig. S9). These examples of Oprm1+ SPNs with strong tuning to action value and trial type were rare, and to better characterize their prevalence we calculated the win-stay versus lose-switch selectivity score for neurons in each trial phase. We found that the D1+, A2A+, and Oprm1+ pathway selectivity scores were similarly distributed (Fig. 5B). In addition, all three pathways showed a similar proportion of neurons significantly tuned to the win-stay and to lose-switch trial type, further supporting a pathway-independent representation of the task (Fig. 5C). The value modulation was often phase-specific and not persistent throughout the trial, and it was common for SPNs to show some value modulation in more than one phase. The proportion of SPNs with win-stay or lose-switch selectivity in each trial phase was similar in all three pathways (Fig. 5D-F).

**FIGURE 5.**
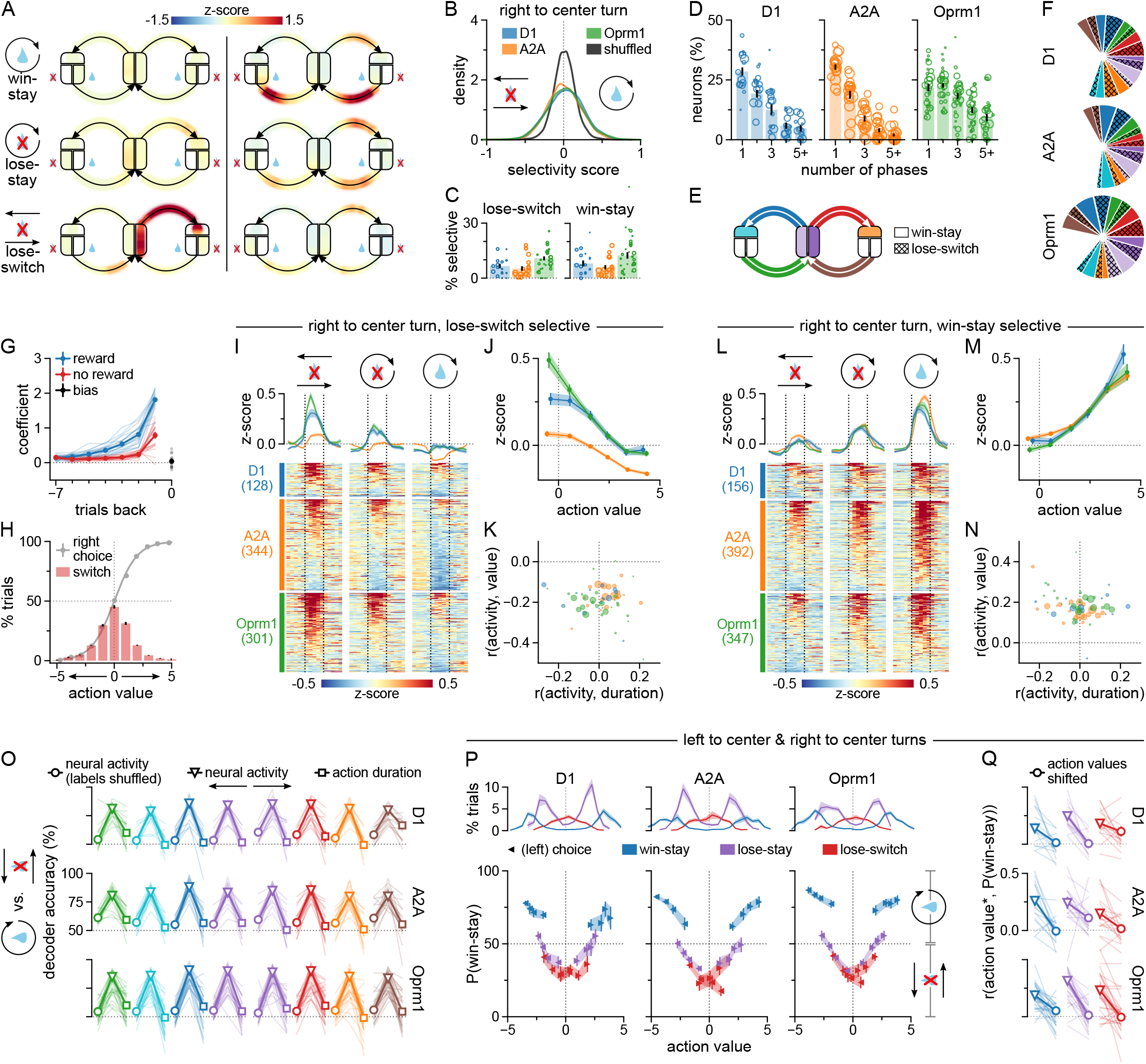
The D1+, A2A+, and Oprm1+ pathways represent all aspects of the choice task structure. A) Example of SPNs with phase-specific evaluation for stay or switch choice. Example Oprm1+ neuron (left side) shows elevated activity in the center port preceding right port choices, as well as during the choice action, only in the context of left-to-right lose-switch trials when the value of the left choice is low. Example Oprm1+ neuron (right side) shows increased activity when returning from the choice port to the center in win-stay trials, when the value of the previous choice is high. Note that it also responds, to a lesser degree, during return movements in lose-stay trials, when the last choice is of intermediate value due to rewards received on trials prior to the recent loss (see G). B) Distributions of the right port to center port turn-specific lose-switch versus win-stay selectivity scores (color code shows pathway). A positive score indicates a neuron’s preferential activation on win-stay, a negative score on lose-switch trials. Selectivity scores: area under the ROC curve, scaled from −1 to 1. For comparison, selectivity scores were calculated using shuffled behavior data (black). (n=1943 D1+ neurons, 6566 A2A+ neurons, 2793 Oprm1+ neurons). C) The weighted mean percentage of neurons significantly selective for lose-switch (left) and win-stay (right) trials during the right port to center port phase (circles: single sessions, radii proportional to the number of neurons, error bars: ±SEM bootstrapped). D) Bar plots of the weighted mean percentage of neurons significantly win-stay or lose-switch selective to several counts of phases. Sessions were weighted by the number of neurons (circles: single sessions, radii proportional to the number of neurons, error bars: ±SEM bootstrapped). E) Illustration of the choice task with phase color-coding used in panels F and O. F) The primary phase-specific win-stay or lose-switch selectivity of SPNs. (n=1943 D1-neurons, 6566 A2A-neurons, 2793 Oprm1-neurons). G) Logistic regression models the impact of the outcomes of the previous seven trials on the current port choice. Mice repeated recently rewarded choices (blue), and were likely to repeat the previous choice even after one reward omission (red). Bias coefficient (black): positive values indicate right port bias. Logistic regression coefficients for individual mice (thin lines) and the mean coefficients (thick lines). H) Average fraction of right port choices (grey dots) and choice switches (red bar plots) as a function of relative action value (binned). Grey curve: logistic regression prediction (data pooled over all mice). Positive (negative) action values predict right (left) port choices. Action values are the sum of coefficients shown in G. Outcome history dictates the coefficient (reward or no reward) used for each of the previous seven trials, side choice its sign (right: positive, left: negative). Dots and bars: mouse average (N=19 mice). Error bars: ±SEM. I) Scaled time raster plots of the average activity of the lose-switch selective neurons in lose-switch, lose-stay and win-stay trials. The dotted lines mark the start and the end of the right port to center port motion. The line plots above the rasters show the population mean activity. J) The mean trial activity of lose-switch selective neurons at various action values (average neuron activity, binned). K) The session-averaged Pearson correlations for lose-switch selective neurons to movement duration versus to action value. Radii reflect the number of selective neurons in the session. L) Scaled time raster plots of the average activity of the lose-switch selective neurons in lose-switch, lose-stay and win-stay trials. The dotted lines mark the start and the end of the right port to center port motion. The line plots above the rasters show the population mean activity. M) The mean trial activity of win-stay selective neurons at various action values (average neuron activity, binned). N) The session-averaged Pearson correlations for win-stay selective neurons. O) Phase-specific decoding accuracy of predicting win-stay versus lose-switch trial type from neuron activity using cross-validated linear SVMs. For each trial phase SVMs were trained on data with the behavior labels shuffled (circles), on the neuron activity data (triangles), or only on phase duration (squares). Accuracy for individual sessions (thin lines) and mean accuracy (thick lines). Sessions weighted by the number of neurons (N=16 D1-sessions, 19 A2A-sessions, 30 Oprm1-sessions). P) The SVMs probability estimates for outcome port (left or right side) to center port turns being win-stay rather than lose-switch as a function of action value. Action value correlates with the SVM prediction confidence. Win-stay trial turns are more confidently predicted in high action value trials. (blue). Lose-switch trial turns are more confidently predicted in low action value trials (red). Lose-stay trial turns with low action value (magenta) appear increasingly like lose-switch turns. Correlation scores are quantified in Q. Binning by quartiles of the pooled action value distributions. Triangles: mean probability estimate of sessions weighted by number of neurons. Shading and error bars: ±SEM (bootstrapped). Top panel: density of each trial type by action value; average of sessions ±SEM. Q) Correlation between action value and the estimated probability of choice port to center turns being win-stay rather than lose-switch (action value sign reversed for left choice trials). Correlation of individual sessions (thin lines) and average correlation of sessions weighted by the number of neurons (thick lines). Color-coding as in P.

To extract the decision variables that accounted for win-stay versus lose-switch, and specifically the impact of reward history on the behavior, we estimated the trial by trial action value using logistic regression (Fig. 5G). The action value estimates accurately reflected the choices and decisions made by the mice (Fig. 5H). To visualize the neuron activity showing trial type selectivity, we focused on a single decision-relevant phase following the outcome in the right port (the right port to center port turn) and the representation of action value in that phase (Fig. 5I-N). We first selected neurons with a significant lose-switch selectivity score in this phase, and plotted the average activity of each neuron during the three main trial types (win-stay, lose-stay, lose-switch).

The lose-switch selective neurons were as expected more active in the lose-switch trial type compared to the win-stay trial type (Fig. 5I). During the lose-stay trial types, these neurons instead showed an intermediary activity profile, suggesting that the population is encoding value rather than trial type identity. We found that the activity of the lose-switch selective neurons was negatively correlated with action value (Fig. 5J). Importantly, since vigor of behavioral response as a metric of motivation could confound the activity signals, we verified that the activity and action value negative correlation could not be explained by a correlation with the action duration (Fig. 5K). We repeated the same analysis for neurons with a significant win-stay selectivity score. The win-stay selective neurons showed increased activity in win-stay trials compared to lose-stay trial in the center to right turn phase, and showed an intermediate activity level in lose-stay trials (Fig. 5L). Correspondingly, the activity of win-stay selective neurons showed positive correlation with the action value, which was not explained by the action duration (Fig. 5M-N).

The action value signal during the return from the right port to center port phase prompted us to determine whether the value and decision variables could be found in other task phases as well. We therefore trained SVMs to predict win-stay versus loses-switch trial types for each of the eight discrete phases of the task. We found that the single neuron activity in the D1+, A2A+, or Oprm1+ pathway contained information to accurately decode the trial type in every trial phase within a session and across sessions (Fig. 5O and Fig. S9). To exclude that the prediction accuracy depended on differences in the structure of the behavior between trial types (e.g. turning speed), we trained SVMs on the action duration and found that they could not accurately predict the trial type, demonstrating that the win-stay or lose-switch activity signals were not reflecting the vigor of the action (Fig. 5O).

The representation of win-stay versus lose-switch we have described could reflect the choice or most recent outcome, rather than the action value. To disentangle these possibilities, we applied the SVM model to the three trials types (win-stay, lose-stay, lose-switch) separately. The SVMs were trained to distinguish lose-switch from win-stay trial types, based on choice or outcome, but not the action value per se. We specifically investigated whether the SVMs would be more confident of the win-stay/lose-switch classification depending on the action value. We found that within each trial type the action value was separately correlated with the confidence in the prediction (Fig 5P-Q). Importantly, this correlation was also found in the lose-stay trial type that was never used for SVM training. This suggests that in addition to choice and outcome, information about the action value is also present in the activity and is used by the SVMs to classify the trial types.

Supporting the evidence for pathway-independent representation of action value and trial type, the SVM decoding of the trial type was similarly dependent on the action value in the D1+, A2A+, and Oprm1+ pathway (Fig. 5P-Q and Fig. S9). To verify that these results were not contingent on the definition of the action value, we calculated action values based on Q-learning and found this model replicated the correlation between SVM confidence and action value (Fig S10). In summary, the action value representation by SPNs was phase-specific during the trial, independent of action duration or motivation, and importantly was evident in all striatal output pathways.

## DISCUSSION

The dorsomedial striatum, and in particular the Oprm1+ striatal patches, are suggested to carry signals on the value of actions, the chosen behavioral strategies, and underlie goal-directed behaviors. We found that the Oprm1+ pathway shows a surprisingly broad tuning to all the investigated behavioral variables, with individual SPNs showing sharp tuning to specific movements, and tuning to discrete actions in a choice task, and therefore do not encode a single decision-making variable such as the type or value of the ongoing trial, and as a population form a continuous representation of the motor and action program that is very similar to the representation in the direct and indirect pathway.

We have shown how the activity of SPNs can describe the entire action space including key variables related to trial type and decision-making (e.g. win-stay strategy, value of the selected action), and importantly that this representation is found in all three major striatal output pathways. We found that the activity did not simply reflect the individual actions as unitary and discrete motions (e.g. left turns) but instead captured the complexity of the task structure, by integrating value and trial phase information for the execution of a specific action. Individual SPN activity tiled the entire task space, forming a continuous representation of all the actions required to perform the task. Supporting the role of these representations for the proper task execution is the conserved and highly stereotypical tuning of individual SPNs to specific actions in a single trial phase over many behavioral sessions and days. Interestingly, the remapping of SPN tuning between the open field and the choice task point to a context-dependent representation of discrete motor-action variables by individual SPNs and further supports that the SPN activity does not represent one dimension of the action (i.e. the concept of a left turn).

It was surprising to find the extent of similarity in the tuning profile between the three molecularly and neuroanatomically distinct output pathways of the striatum, considering the suggested specialization of each pathway in motor programs. The standard basal ganglia model of antagonistic signals in the direct versus indirect pathways is supported by optogenetic manipulations, demonstrating their opposing or differential effects on reinforcement, choice, and action sequences (Geddes et al., 2018; Kravitz et al., 2012; Tai et al., 2012). In contrast, recording of the striatal pathway activities during motor behavior does not reveal a clear distinction between the pathway activities (Cui et al., 2013; Tecuapetla et al., 2016). Interestingly, the encoding of action value has been found in dorsomedial as well as the more motor-related dorsolateral striatum (Stalnaker et al., 2010), including a pathway bias in value encoding (Shin et al., 2018), supporting a complex and rich representation of motor and decision-making variables across striatal regions.

The temporally organized activation of SPN subtypes has been proposed to be a key mechanism in representing the motor program either as start-stop signals (Jin and Costa, 2010) and action chunking (Graybiel, 1998) or as continuous sequence representations (Akhlaghpour et al., 2016; Geddes et al., 2018; Sales-Carbonell et al., 2018), while other studies have emphasized role of SPNs in categorical coding of value (Samejima et al., 2005; Wang et al., 2013). The activity in dorsolateral striatum during a simple locomotion task could not be clustered (Sales-Carbonell et al., 2018), suggesting that the motor-related activity of SPNs is high-dimensional and instead contains mixed representations of several variables. Studies have shown that the dorsal striatum contains neurons that encode spatial information (Hinman et al., 2019; van der Meer et al., 2010), visual and tactile information (Reig and Silberberg, 2014), delay periods (Akhlaghpour et al., 2016) and time(Mello et al., 2015), signals that together can shape the representation of behaviorally relevant information. The diverse representation of for example space, time and trajectory that previously have been found in striatal neurons suggest a complex integration of various behaviorally relevant signals in individual SPNs. Our findings support a model where the encoding of action sequences, strategy and value are simultaneously represented during action selection, by integrating motor and decision signals that are uniquely representing a behavioral context, to ultimately form a continuous representation that is relevant for proper task learning and performance. Models of the basal ganglia as an actor-critic system suggest that the patch compartment plays the role of the critic while the actor is implemented in the matrix compartment (Barto, 1995; Doya, 1999). Even if our findings on the clear overlap between representations found in Oprm1+ as well as D1+ and A2A+ pathways is challenging this view, it is interesting to note that recent work in deep reinforcement learning has proposed that it could be beneficial to share parameters between actor and critic in early layers of artificial neural networks (Mnih et al., 2016). Therefore, a similar neural representation in patch and matrix pathways might analogously form a basis for both evaluative and selective processing in the downstream targets.

It will be valuable to determine the role of the different input pathways and to what extent they shape the rich representation found in the striatal pathways. The dorsomedial striatum receives prominent inputs from the frontal cortex, which contains key decision signals (Hwang et al., 2019; Padoa-Schioppa and Assad, 2006). The multi-tuning and multiplexing of task-relevant signals has been observed in a number of cortical circuits (Musall et al., 2019; Steinmetz et al., 2019; Stringer et al., 2019), and some of these circuit calculations are likely to be transmitted and represented in the striatal circuitry (Peters et al., 2019). Even if the corticostriatal functional connectivity has not been comprehensively defined at the detail of single SPNs, the corticostriatal organization has been proposed to generally be pathway-specific (Gerfen, 1989; Lei et al., 2004; Wall et al., 2013), although evidence also points to converging organization (Smith et al., 2016; Zheng and Wilson, 2002). This organization of corticostriatal inputs carrying signals that are distributed to all types of SPNs could underlie the broad tuning we have observed.

The technical limitations in terms of the dynamics captured by calcium imaging could obscure some aspects of differential activity in the three output pathways, such as pathway-specific tonic firing rate changes or differences in event-locked latency, and our conclusions therefore are based on a general description of how the observed SPN activity reflects the main aspects of behavior. In addition, more detailed analysis of the kinematics and recording of an even larger number of SPNs or simultaneous recording of SPNs in different striatal regions can provide an even better description of the differences and similarities in the signals represented by different striatal outputs.

We found that Oprm1+ SPNs integrate detailed aspects of motor-action signals, carrying signals about the movement, behavioral context, trial phase, and strategy, to produce a rich representation of the progression through the task as well as action value and choice variables, and that this complete representation is found in the D1+ direct and A2A+ indirect striatal pathways as well. The high-dimensional representation of the entire task and the context-dependent tuning of individual SPNs must be considered when developing circuit models of the basal ganglia to understand motor and action behavior. We therefore propose that the dorsomedial striatum broadcasts an ergocentric (greek [έργον]: work, task) representation of the entire task to all downstream targets: the neuron activity is a merged representation of the phase-specific action together with abstract representation of key decision-making variables of the task space including upcoming choices and the value of specific actions.

## AUTHOR CONTRIBUTIONS

M.W. performed behavior experiments, analyzed and interpreted data. E.W. analyzed and interpreted data. I.L. interpreted data and performed surgeries and behavior experiments. O.T. characterized the Oprm1-Cre mouse line. K.M. designed the study, interpreted data, and wrote the manuscript. All authors discussed and commented on the manuscript.

## ACKNOWLEDGEMNTS

Funding for this study was provided by the Swedish Research Council (Vetenskapsrådet, Medicin och hälsa), the Swedish Brain Foundation (Hjärnfonden), Karolinska Institutet (KID doctoral funding for M.W., E.W., O.T.,).

## METHODS

### RESOURCE AVAILABILITY

Further information and requests for resources and reagents should be directed to and will be fulfilled by Konstantinos Meletis.

### Materials Availability

No unique materials were produced in this study.

### Data and Code Availability

Data and Python code to reproduce all figures in this work will be made freely available online upon publication.

## EXPERIMENTAL MODEL AND SUBJECT DETAILS

The 19 adult female and male transgenic mice (25–35 g) used in experiments were kept on a 12-hour light/dark cycle. We used D1-Cre (Drd1-cre EY262Gsat), A2A-Cre (Adora2a-cre KG139Gsat), and Oprm1-Cre mice (Märtin et al., 2019). Mice were single-housed after the surgeries and placed on food restriction during behavioral training and recording in the 2-choice task (maintained at min. 85% of their free-feeding body weight). All procedures were approved by the Swedish local ethics committee for animal experiments (Stockholms djurförsöksetiska nämnd, approval N166/15).

## METHOD DETAILS

### Surgeries

For stereotactic surgery, the mice were anesthetized with isoflurane (2% in air) and administered buprenorphine (0.03 mg/kg, subcutaneous) for analgesia. Buprenorphine was also administered for post-surgery pain-relief. A feedback-controlled heating pad maintained body temperatures at 36°C throughout the surgical procedures. All animals were unilaterally microinjected with 400 nl of AAV5-CAG-Flex-GCaMP6s into the dorsomedial striatum in the right hemisphere (AP: 1.0, ML: 1.25, DV: −2.3; rate of 100 nl/min). The pipette was retracted 5 minutes after the injection finished. Two weeks after the viral injection, gradient-index (GRIN) endoscope lenses (Inscopix) of a 1 mm diameter were implanted 100-200 μm above the viral injection site. Prior to lowering the lens, portions of the overlaying cortex were aspirated using a 0.5 mm-diameter, blunt-point needle with sharpened edges which was attached to a vacuum pump. The GRIN lens was fixed in place using dental cement. 4-6 weeks after GRIN lens implantation, the baseplate was anchored to the skull with dental cement to support the detachable miniscope (Inscopix) during imaging. To determine the optimal placement of the baseplate, the field of view was monitored live throughout the procedure using the baseplate-attached miniscope. The miniscope was not refocused over the course of the experiments in order to improve tracking of individual neurons across days.

### Histology

At the end of the experimental procedure mice were deeply anaesthetized with pentobarbital and then transcardially perfused with 0.1 M PBS followed by 4% paraformaldehyde in 0.1 M PBS. Brains were removed and post-fixed in 4% paraformaldehyde overnight at 4°C and then washed and stored in 0.1 M PBS. Coronal 80μm sections were cut using a vibratome (Leica VT1000, Leica Microsystems, Nussloch GmbH, Germany). Immunostaining was performed on free-floating sections in glass wells. Sections were incubated for 1 hour in 0.3% TritonX-100 in Tris–buffered saline (38mM Tris-HCl, 8mM Trizma base, 120mM NaCl in extra pure water) and treated with a preheated (40°C) antigen retrieval solution (10mM sodium citrate, 0.05% Tween20, pH:6) for 1-2 minutes. In order to block non-specific antibody binding, sections were incubated in 5% Normal Donkey Serum in TBST (0.3% TritonX-100 in Tris– buffered saline) for 1 hour at room temperature. Sections were subsequently incubated overnight with primary antibodies at room temperature. The day after, sections were washed twice for 10 minutes in TBST, then incubated with secondary antibodies for 4 hours at room temperature, and finally imaged. Primary antibodies used: goat anti-GFP (1:1000 dilution, abcam: ab5450); rabbit anti-Tyrosine Hydroxylase (1:100 dilution, abcam: ab112). Fluorophores of secondary antibodies: Alexa Fluor-488 and Cy3 from Jackson ImmunoResearch Laboratories. Assessment of GRIN lens placement was based on the lesion from the lens in the tissue. Animals with misplacement of GRIN lens were excluded from the study.

### Open field task

We tracked the mice in the open field (49×49 cm) using DeepLabCut (DLC) (Mathis et al., 2018). We manually labeled the base of the tail, the center of the body and the left and right ears in 1000 frames, sampled from all sessions. After running DLC on every open field video, we transformed the video coordinates (in pixels) to world coordinates (in cm) using a perspective transform matching the four corners of the box.

We based our movement analysis on two markers tracked by DLC: the base of the tail and the center of the body. We used a custom particle filter (2000 particles; diagonal gaussian observation noise with σ=1 cm; diagonal gaussian innovation noise with σ=1 cm/frame; gaussian likelihood penalty on distance between tail base and center of body with μ=3 cm and σ=1 cm; *systematic resampling* as defined in (Doucet and Johansen, 2011) to estimate smoothed x- and y-position as well as running speed, allocentric body direction, angular speed and body elongation.

We then developed a custom optimization algorithm to classify each video frame as either *left turn, right turn, running* or *stationary*. First, we chose a scoring function *P*_*b*_*(s,e)* that defined how well a segment starting at frame *s* and ending at frame *e* would fit behavior *b*. In particular, we chose:

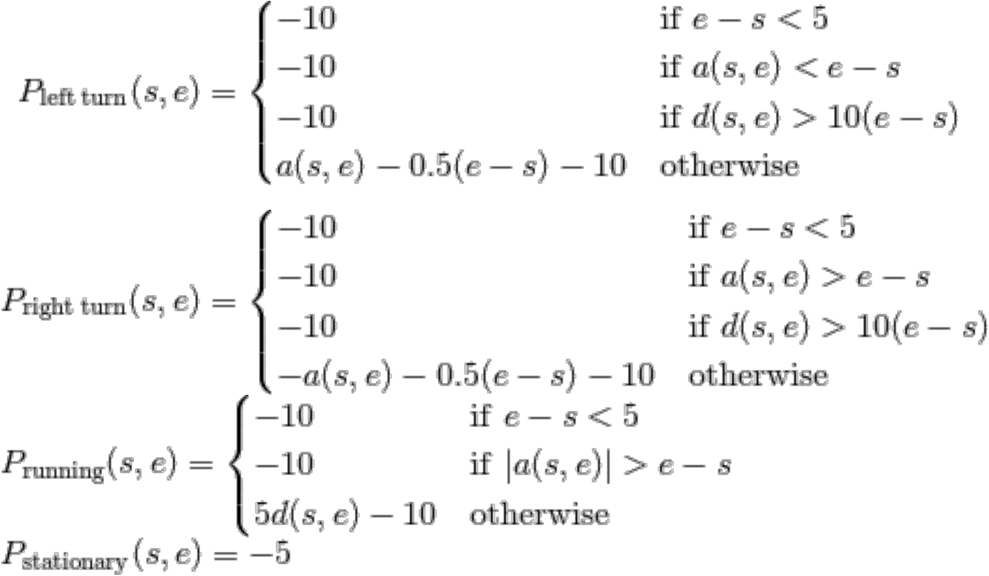

where *a(s, e)* is the signed angle difference between the animal in frame s and frame *e* measured in degrees, and *d(s, e)* is the distance in cm between the animal’s position in frame *s* and frame *e*. Note that both *a(s, e)* and *d(s, e)* can be computed in constant time. The negative terms acted as a cost on the number of segments to encourage the segments to be as long as possible. Having this definition, we wanted to find segments of behaviors such that the total score was maximized, i.e.

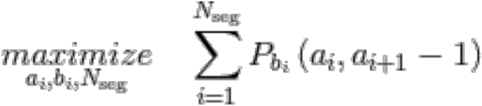

where *N*_*seg*_ is the number of segments, *a*_*i*_ is the start of segment *i* and *b*_*i*_ is the behavior of segment *i* (*a*_*1*_*=1, a*_*i*_*<a*_i+1_, *a*_*N*_=number of frames). The optimal *{N*_*seg*_,*a*_*i*_,*b*_*i*_*}* can be found very efficiently with dynamic programming on the recurrence relation:

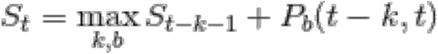

To improve running time further, we constrained *k*<2000 so that the algorithm ran in linear time.

### 2-choice task

We trained the food-restricted mice to perform a self-paced, probabilistic 2-choice task inside custom-built operant chambers which contained three nose poke ports. The mice initiated a choice in the center port, and were then free to choose one of the two side ports. One side port yielded sucrose solution with 75% probability upon entry (15% sucrose, 3.75 μl), while the other yielded nothing. After a reward, there was a 5% probability of a reward port switch. When no reward was delivered (due to failed trial initiation, incorrect port choice or reward omission) the rewarded side remained the same. There were no cues to indicate the correct port choice nor the occurrence of a reward port switch. The volume of the sucrose reward was doubled in a random 10% of the rewarded trials. LEDs located in the ports indicated whether a trial was in the initiation phase (center LED on) or in the choice phase (side LEDs on). A 350 ms infrared beam break was required to trigger any port. The task was controlled and task events recorded using a pyboard microcontroller.

Animals failed to trigger the reward ports in almost half of their attempts, predominantly due to having failed to wait out the delay in the center port first, exiting it early (<350ms; see Figure 2C, delay error). This suggests they disregarded the LED cues indicating whether the initiation nose poke was successful and remained unaware of their error. Thus, we included every side port entry in the analysis, irrespective of whether or not the choice was properly initiated in the center port.

The effect of trial outcome history on port choice, used to compute trial-by-trial relative action value, was estimated by logistic regressions, as reported previously (Tai et al., 2012). The following regression model was fitted for each animal separately:

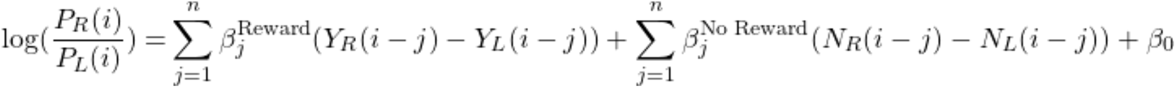

P_R_(i) represents the probability of choosing the right side port on the current side port entry, indexed i. Y_R_(i−j) *∈ {0,1}* and N_R_(i−j) *∈ {0,1}* denote whether or not a right port choice was rewarded or unrewarded j entries back, respectively. P_L_(i), Y_L_(i), and N_L_(i) code the equivalent variables for left port choices. The coefficients β_j_^Reward^ and β_j_^No Reward^ hence capture the effects of obtaining or not obtaining a reward j trials ago on the current choice. The intercept term β_0_ subsumes any static bias towards one port or the other. n, the number of past trials included in the regression, was set to 7. The coefficients were fit using maximum likelihood. The model-predicted trial-by-trial log-odds of side-port choice, negative values favoring left choice, positive right choice, served as relative action values.

To verify that our results were not dependent on calculating action values using logistic regression we also calculated action values based on Q-learning (Figure S10). We let *Q*_*R*_*(t)* denote the value of choosing the right port and *Q*_*L*_*(t)* the value of choosing the left port. Assuming the animal enters port *p ∈ {L,R}* at time *t*, we update the corresponding value

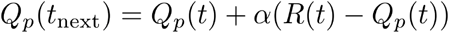

where α the learning rate and *R(t) ∈ {0, 1}* is the outcome. Given Q_R_ and Q_L_, we used a logistic function to estimate the probability of the animal choosing the right port

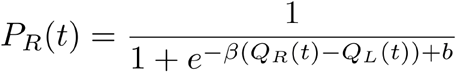

where *β* is controlling slope of the function, and therefore the explore-exploit trade-off, and *b* is capturing any static bias towards either side. We fitted the three parameters *α, β* and *b* separately for each animal by maximizing the likelihood (equivalent to minimizing the negative log-likelihood)

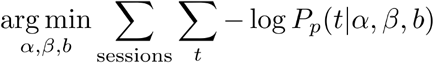

where *p ∈ {L,R}* is the choice at time *t* and *P*_*L*_ *= 1 - P*_*R*_. We used the downhill simplex algorithm (scipy.optimize.fmin) to minimize the negative log-likelihood.

### Imaging data acquisition and preprocessing

Raw calcium imaging videos (1440×1080, 20 fps) were acquired using miniscopes and the nVista Acquisition Software (2.0.4, Inscopix). To accurately align calcium transients and behavior, we recorded the sync pulses provided by the miniscope’s data acquisition box in the output of the microcontroller controlling the operant task. The sync pulses also served to trigger a Blackfly USB3 video camera (FLIR) used to record both the operant and the open field experiments. 2-choice task imaging sessions lasted approximately 1 hour, open field sessions 20 min (see Supplementary tables S1-S3). Calcium imaging videos were spatially downsampled (2×2 pixel bins), cropped, and motion corrected using the Inscopix Data Processing Software (1.2.0) and exported as Neurodata Without Borders (NWB) files. We used the CaImAn (1.4.2) (Giovannucci et al., 2019) implementation of the CNMF-E algorithm (Zhou et al., 2018) to extract spatial filters (i.e. ROIs) and deconvolved fluorescence traces of individual neurons. Imaging frames were downsampled (2×2 pixel bins) once more prior to running the algorithm. To adjust for bleaching, the rolling z-score (±5 min window) of the deconvolved activity traces was computed and used in all subsequent analysis. To match individual neurons across recording sessions, we used CaImAn’s ROI-registration function. Prior to analysis, we excluded sessions with data alignment issues (dropped imaging frames) or during which the mouse did not engage in the task.

### Definition of trial phases

Each calcium imaging frame was assigned to a trial phase solely based on the infrared beam brakes detected around that frame. Frames recorded while a particular port’s beam was broken were registered as occupations of that port (i.e. nose pokes). When no beam was broken, frames were classified as movements between the port occupied last and the port entered next. We excluded thus labeled movements bypassing the center port (side port to side port), movements lasting longer than 1.5s (animal unengaged), as well as movements starting and ending in the same port (reentries), from further analysis. Nose pokes preceding unengaged periods or reentries were also excluded. We divided center port nose pokes into two different phases, depending on which side port was entered next (upcoming left or right choice). Side port nose pokes were split into three phases: the *delay* phase preceding the outcome presentation (<350ms in port) and the two mutually exclusive outcome phases (>350ms in port), *omission* and *reward*. Note that this classification approach disregards whether ports were triggered or not, i.e. nose pokes shorter than 350 ms and nose pokes addressing inactive ports (due to a delay error in the previous nose poke) are included in the analysis.

For drawing trial heatmaps (e.g. Figure 3B) and for sub-phase decoding (Figure 3L-N), we assigned a *progress score* to each frame, defined as the count of consecutive previous frames spent in the current phase, divided by the total number of frames in the current phase. Note that this scaling of the progress was therefore based on *time* between beam breaks, not on physical space.

### Pooled-population activity trajectories

The activity trajectories in figure 2H are based on single-trial activity pooled over neurons of all Cre lines and recording sessions (without matching neurons across sessions). Individual trials in which an animal did not follow the task structure perfectly, e.g. by reentering a port or disengaging from the task mid-way, were excluded from this analysis, as were trials in which any phase lasted fewer than 250 ms (5 frames). Whole sessions were excluded if there were fewer than 20 trials left of any of the three trial types visualized (right win-stay, right lose-stay and right lose-switch). To pool the single trial data from different sessions into pooled-population pseudo-trials, we contracted all trials to uniform length by binning the trial phases in scaled time (4 bins per phase, resulting in a total of 20 data points per trial). For every trial type, we then randomly sampled trials of that type from every neuron in every session, combining them into pseudo-trials of pooled-population activity. Trials were drawn without replacement; the total number of pseudo trials of a specific type was therefore determined by the minimum number of trials of that type performed in any session. We applied PCA to one pooled data set obtained in this way, treating the neurons as features to be reduced and the time points of the concatenated pseudo-trials as samples. We then plotted the average trial trajectory using the first 3 principal component scores (thick lines in 2H). To evaluate the reproducibility of this approach, we resampled the pooled-population data set several times, obtaining different subsets and combinations of single-trial data, and projected these pseudo-trials into the same PCA space (thin lines).

Note that the random drawing of trials forced the neurons in the pseudo-session to be conditionally independent given the trial phase and type, and therefore effectively canceled out any principal components that were not aligned to our parameterization of the task. This was done on purpose to enhance the task-relevant structure in the neural activity and we term the principal components calculated this way “task-related principal components”.

To test whether the activity in general was low-dimensional, we furthermore applied PCA directly to the deconvolved and z-scored traces of each session separately (Supplementary Figure S5B-D). Additionally, to estimate whether the activity was confined to a non-linear manifold, we calculated the *internal dimensionality* (Rubin et al., 2019): for a manifold of (non-linear) dimension *d*, we expect the average number of neighbours within a small L2-distance *r* to be proportional to *r*^*d*^ (volume of a hypersphere). To estimate *d*, for each point in time in each session, we found the 500 other timepoints with most similar activity (L2 distance). Using these 500 points, we first discarded the 3 closest ones, and then linearly regressed the logarithm of the L2-distance to the logarithm of the number of points within that distance. We performed one such regression for each point in time (50ms) in each session. Our estimate of the dimensionality of the activity during a session was the averaged linear coefficients from all the regressions (Supplementary Figure S5E).

### Visualization of trial heatmap

To create the trial heatmaps we first assigned a trial phase and a phase progress (0 to 1) to each calcium imaging frame. Next, we used these to calculate a pixel coordinate in the schematic (full image was 501×251 pixels). For the center port, the y-coordinate increased with increased phase progress, such that 0 was at the bottom and 1 was at the top of the rectangle. Two separate x-coordinates were used depending on the upcoming choice. For the side ports, the y-axis was reversed compared to the center port so that increasing phase progress was going downwards. The scaling was such that the beginning of the delay phase always was at the top and the end of the delay phase was at the indicated line. The reward and omission phases similarly progress downwards. The omission phase is usually shorter than the reward phase (i.e. the animal exits the port earlier), but scaled to have the same length in the visualization.

For leftwards movements, i.e. center-to-left and right-to-center, the x-coordinate was linearly decreasing with phase progress. For rightward movements, i.e. center-to-right and left-to-center, it was linearly increasing with phase progress. The y-coordinate of the four movements was the square of the progress, followed by a translation and scaling. The phase progress therefore does not follow the tangent of the parabola between the respective ports but rather the x-coordinate.

We then calculated the average deconvolved z-scored activity for each pixel and smoothed the averages with a gaussian kernel (σ = 7 pixels). Finally, we applied the colormap to the smoothed value at each pixel. We set the color according to the mean activity and the transparency according to how many times the (smoothed) pixel was visited.

### Calculation of neuron tuning scores

We created surrogate data by first dividing the session into blocks of consecutive left and right trials and then shuffling these blocks. This procedure meant most of the behavioral statistics of the session were preserved (number of left and right trials, stay and switch probabilities, time between trials and between phases within a trial, etc), while the connection to the neural activity was broken.

To calculate the tuning score, we calculated the average z-scored fluorescence during each of the 12 trial phases. As a comparison, we did the same for 1000 iterations of blocks shuffled as described above. We classified a neuron as *positively tuned* to a trial phase if the average activity using the real data was larger than in 99.5% of the shuffled iterations or, reversely, as *negatively tuned* if it was less than 99.5% of the shuffled iterations. In addition to these discrete classifications, we also defined a *tuning score* as the real average minus the grand average of the shuffled iterations and divided by the standard deviation of the averages of the shuffled iterations. We call the 12 tuning scores for a given neuron the *tuning profile* of that neuron.

For the open field, we used a similar procedure to define positively and negatively tuned neurons, as well as tuning scores, to the four behaviors. The shuffle was taken over the behavioral segments (left turn, right turn, running, stationary) without consideration of transition probabilities.

To determine how well a neuron’s activity discriminated win-stay from lose-switch trials in the choice task, we computed a *selectivity score* based on receiver operating characteristic (ROC) analysis. The selectivity score was defined as the area under the ROC curve (AUC), scaled from −1 to 1, i.e. equaled 2 x (AUC - 0.5). To establish whether a neuron’s selectivity score was statistically significant, we opted for a similar approach as described for the phase tuning scores above. For every neuron, we computed 1000 selectivity scores based on shuffled behavior data for comparison with the actual selectivity score. If the actual score was higher than 99.5% of the shuffled data scores, we classified the neuron as significantly win-stay tuned, if instead it was lower than 99.5% of the latter, we designated it significantly lose-switch tuned.

To visualize any functional clustering of the tuning to the twelve phases, we stacked the 12-dimensional tuning profiles of all neurons from all pathways. We reduced the dimensionality to two using tSNE (perplexity 30; 10000 iterations; PCA initiation) (Supplementary Figure S5F). To quantify the clusteredness of the tuning profiles, we calculated the silhouette coefficient (Rousseeuw, 1987) assuming clustering by pathway (Supplementary Figure S5G), by the index of the strongest tuning (primary tuning; Supplementary Figure S5H), or by k-means clustering (Supplementary Figure S5I).

To see how trial type (value) influenced the clusteredness of the tuning profiles, we calculated the mean activity for each neuron to each trial phase in each trial type (12*3 = 36 dimensions), and ran agglomerative clustering on this combined data (Supplementary Figure S5I).

### Support vector machines to decode behavior from neuron activity

We z-scored the deconvolved fluorescence traces for each neuron. Then, for every phase of each individual trial, we calculated the average z-scored deconvolved fluorescence for all the neurons. Using the trial phase as the label with the vector of average fluorescences as corresponding features, we trained support vector machines (SVMs) with linear kernels and slack *C*=1. To handle multiclass labels (12 different trial phases), we employed the *one-versus-one* strategy.

In Figure 3J, we randomly selected a subset of neurons and randomly divided 80% of the trial phases into a training set and 20% into a test set. We repeated this procedure 10 times for each session and number of selected neurons (steps of 5 neurons) and reported the average accuracy of these 10 iterations. Accuracy was measured as the fraction of trial phases correctly predicted without accounting for the frequency of each label. Note that this is not traditional 5-fold cross-validation because the train and test split (as well as the selection of neurons) is redrawn in each iteration. In the open field (Figure 1O) we used a similar procedure, but with segments of behavior rather than trial phases.

In Figure 3K, we did five iterations of 80%-20% train-test splits for each session and used all the available neurons in each iteration. We calculated the fraction of each phase type correctly recalled for each session and averaged these values weighted by the number of neurons in the respective session. In Figure 1P we used the same principle to calculate confusion matrices.

When decoding across days (Figure 4E) we only included matched neurons. For all pairs of sessions from the same animal, we trained five iterations of support vector machines on 80% of the trial phases from the session with the earliest recording date. We tested the SVM from each iteration on the remaining 20% of the same session (*same-day-decoding*) as well as on all the trial phases from the other session (*across-days-decoding*). We calculated the average accuracy for each pair of sessions. In Figure 4E, each line corresponds to one such *pair*. Note that this implies the same session is included multiple times. For each genotype and bin of days, the thick lines indicate the average accuracy over all pairs, weighted by the number of matched neurons.

For decoding lose-switch versus win-stay, we averaged the deconvolved and z-scored traces for each phase in each trial, as described above. We then discarded side-port phases and analyzed the remaining six trial phases separately. For each type of trial phase, we discarded all win-switch or lose-stay trials and trained separate SVMs to predict whether the individual phases belonged to a win-stay or a lose-switch trial. Accuracies are reported as the fraction of trial phases that were correctly predicted in a 5-fold cross-validation schema. For comparison, for each decoding we also repeated the respective procedure with one instance of shuffled data created with the same algorithm as when calculating the tuning scores. For Figure 5O, we also compared each SVM trained on neural data to an SVM trained on a single feature, namely the duration of the trial phase that is to be predicted. If the difference in neural activity was only due to different vigor of the movements, these SVMs would reach the same or better performance as the ones trained on neural data.

To see how the SVM decoding varied with the action value, we took the SVMs trained on lose-switch and win-stay trials applied them to all three types of trials. Note that these SVMs never saw any lose-stay trials in the training phase and therefore can never predict lose-stay. Instead of using the binary classification labels, we used the probability they assigned to win-stay over lose-switch for each trial. In Fig. 5P, we binned trials by trial type and action value (quartile bins) and showed the mean probability outputted by all SVMs for the respective bin. We also correlated this probability (without binning) with the action value of each trial separately for each trial type. For comparison, we calculated the same correlation on data where the action value was either shifted forward or backward 10-30 trials (with wrap-around). This preserved the trial-by-trial structure in the action value but dissociated it with the actual neural activity.

## QUANTIFICATION AND STATISTICAL ANALYSIS

Statistically, this work was exploratory rather than confirmatory; there were no formal a-priori hypotheses. Therefore, no statistical methods could be used to determine sample size. For the same reason, we have also promoted descriptive over inferential statistics and in particular refrained from reporting p-values between groups.

All animals were recorded in multiple sessions. Unless stated otherwise, we did not match ROIs across sessions, treating neural traces recorded in different sessions as originating from independent units/neurons. Thus, in figures where the units of measure are neurons (e.g. tuning distributions), the same neuron may be included multiple times. In figures where the units of measure are sessions (e.g. decoding), we calculated averages weighing sessions by the number of recorded neurons, thereby emphasizing sessions with many neurons over those with few. We used the same weighing when bootstrapping SEMs. Supplementary Tables S1-S3 show all the sessions included in the analysis, as well as the number of detected neurons and performed trials.

### Analysis software

In addition to the explicitly mentioned tools, we used Python 3.7, with numpy 1.17.3 (Walt et al., 2011), scipy 1.3.0 (Virtanen et al., 2020), pandas 0.25.3 (McKinney, 2010) throughout. The support vector machines and other machine learning tools were from scikit-learn 0.21.3 (Pedregosa et al., 2011). Some computationally heavy operations were implemented in Cython (Behnel et al., 2011). Plots were rendered with the matplotlib 3.1.1 (Hunter, 2007), seaborn 0.9.0 and figurefirst 0.0.6 (Lindsay et al., 2017) plotting libraries. Figures were edited in Inkscape (0.92.4).

## Supplemental Video titles and legends

**Supplementary video 1**. Supplementary to Figure 1. Video shows two minutes of open field behavior overlayed with DeepLabCut tracking markers and segmentation labels.

**Supplementary video 2**. Supplementary to Figure 3. Video illustrates phase tuning of the example neurons.

**Supplementary video 3**. Supplementary to Figure 5. Video example showing phase-by-phase, cross-validated SVM predictions of trial phase and trial type.

## RESOURCES TABLE

| REAGENT or RESOURCE | SOURCE | IDENTIFIER |
| --- | --- | --- |
| **Antibodies** |  |  |
| Goat anti-GFP | Abcam | ab5450 |
| Rabbit anti-Tyrosine Hydroxylase | Abcam | ab112 |
| Donkey anti-rabbit Cy3 | Jackson ImmunoResearch Laboratories | 711-165-152 |
| Donkey anti-goat Alexa Fluor-488 | Jackson ImmunoResearch Laboratories | 705-545-003 |
| **Bacterial and Virus Strains** |  |  |
| AAV5-CAG-Flex-GCaMP6s | Addgene | 100842-AAV5 |
| **Experimental Models: Organisms/Strains** |  |  |
| Mouse: D1-Cre (Drd1-cre EY262Gsat) | Jackson Laboratories | EY262Gsat |
| Mouse: A2A-Cre (Adora2a-cre KG139Gsat) | Jackson Laboratories | KG139Gsat |
| Mouse: Oprm1-Cre | Meletis Lab | Oprm1-Cre |
| **Software and Algorithms** |  |  |
| Python | www.python.org | 3.7 |
| Numpy | (Walt et al., 2011) | 1.17.3 |
| Scipy | (Virtanen et al., 2020) | 1.3.0 |
| Pandas | (McKinney, 2010) | 0.25.3 |
| Scikit-learn | (Pedregosa et al., 2011) | 0.21.3 |
| Cython | (Behnel et al., 2011) |  |
| Matplotlib | (Hunter, 2007) | 3.1.1 |
| Seaborn | www.seaborn.pydata.org | 0.9.0 |
| Statsmodels | (Seabold and Perktold, 2010) |  |
| FigureFirst | (Lindsay et al., 2017) | 0.0.6 |
| CalmAn | (Giovannucci et al., 2019) | 1.4.2 |
| Inscopix Data Processing Software | Inscopix | 1.2.0 |
| Inkscape | www.inkscape.org | 0.92.4 |
| Custom python and cython scripts | This paper |  |

