## Supplementary Figures and Data for "Complete representation of action space and value in all striatal pathways"

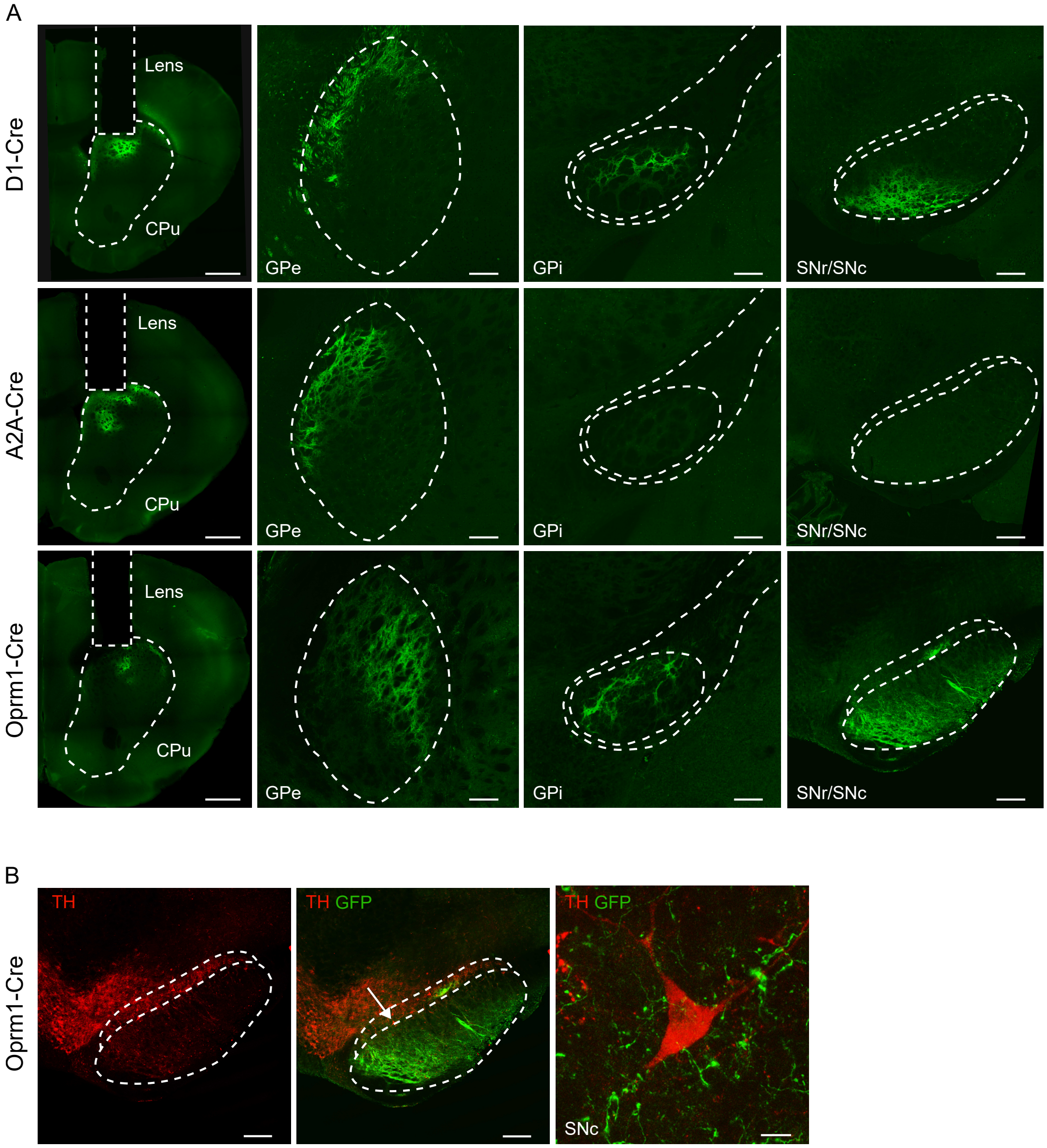


**Figure S1: Supplementary to Figure 1.** A) Higher resolution version of Figure 1B. Images show Cre-dependent GCaMP6s expression (green) in caudate putamen (CPu), globus pallidus externa (GPe), globus pallidus interna (GPi), and substantia nigra (pars reticulata and compacta, SNr/SNc). Scale bars: 1mm for leftmost column, 200$\mu$m otherwise. B) Images show substantia nigra pars reticulata and compacta (SNc) stained for tyrosine hydroxylase (TH, red). Scale bars: 10$\mu$m for rightmost image, 200$\mu$m otherwise.

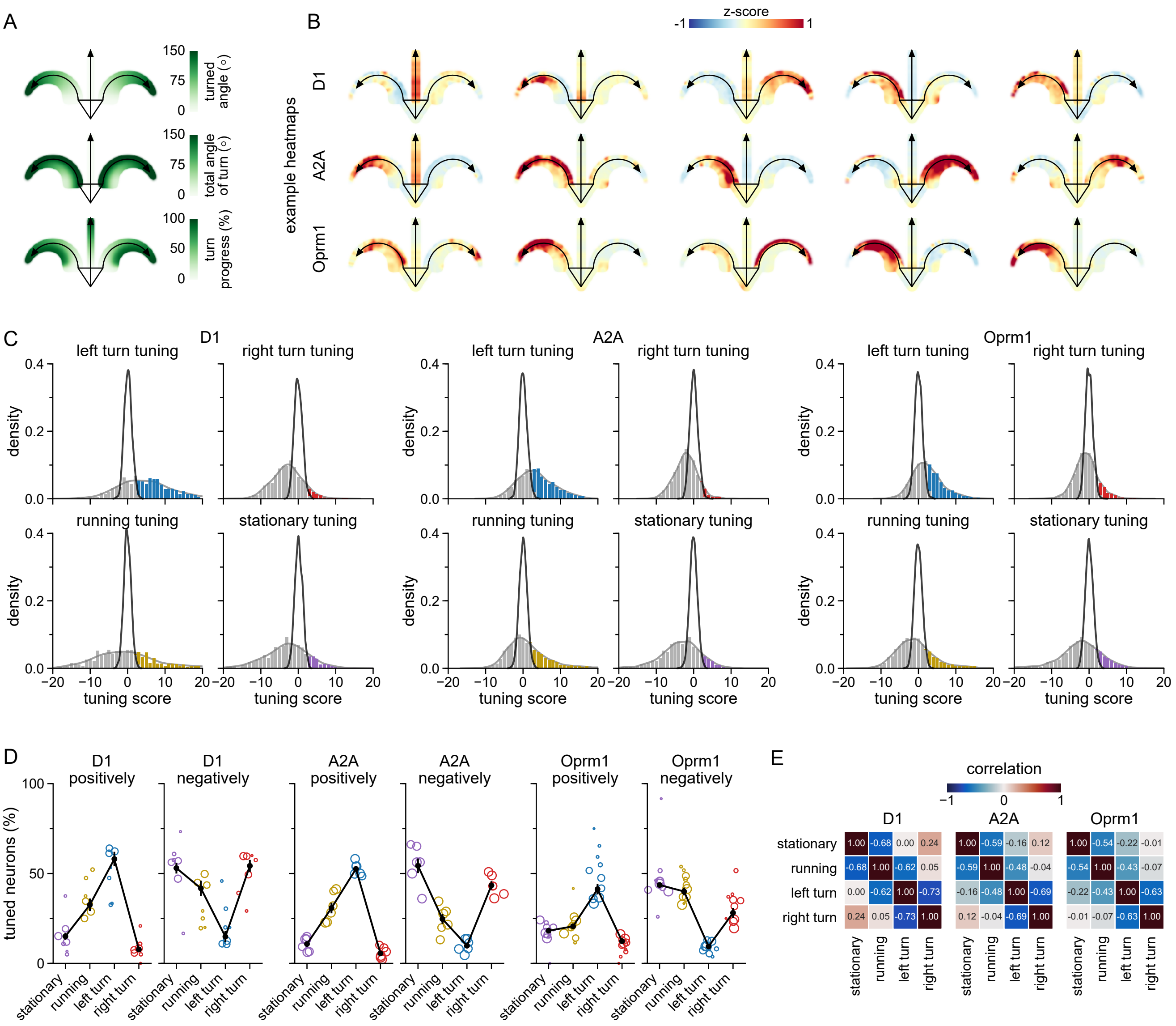


**Figure S2: Supplementary to Figure 1.** A) Legend for the heatmaps. The angle along the curved arrows follows the turn angle of the animal (top). The radial component indicate the total length of the turn, with long turns at the top and short turns at the bottom (middle). As a consequence of these rules, the scaled time of the turn is as indicated bottom panel. B) Five example neurons of each type. Note that different neurons respond to different parts of the turn. C) Distribution of tuning scores per Cre-line and behavior. Black line indicate shuffled distribution, colored bars indicate cells *positively* tuned to the respective behavior. D) Fraction of neurons positively (increased firing) and negatively (decreased firing) tuned to running bouts, turns and phases of neither (*stationary*). E) Correlations between the tuning scores to different behaviors.

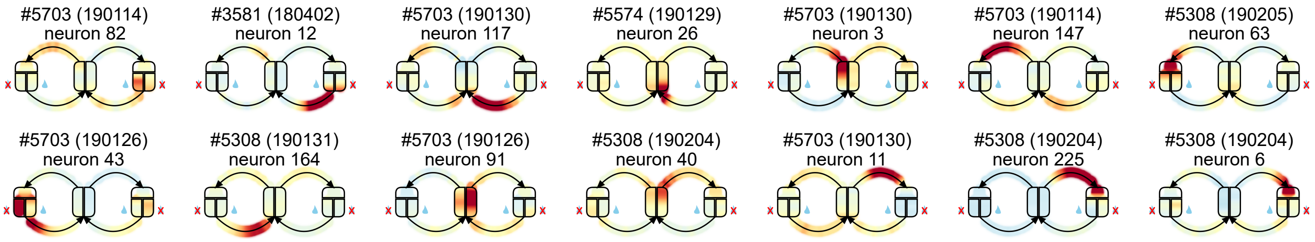


**Figure S3: Supplementary to Figure 3.** Example of Oprm1+ neurons responding to different parts of the task. Panel titles indicate animal ID, date of recording (YYMMDD) and neuron ID.

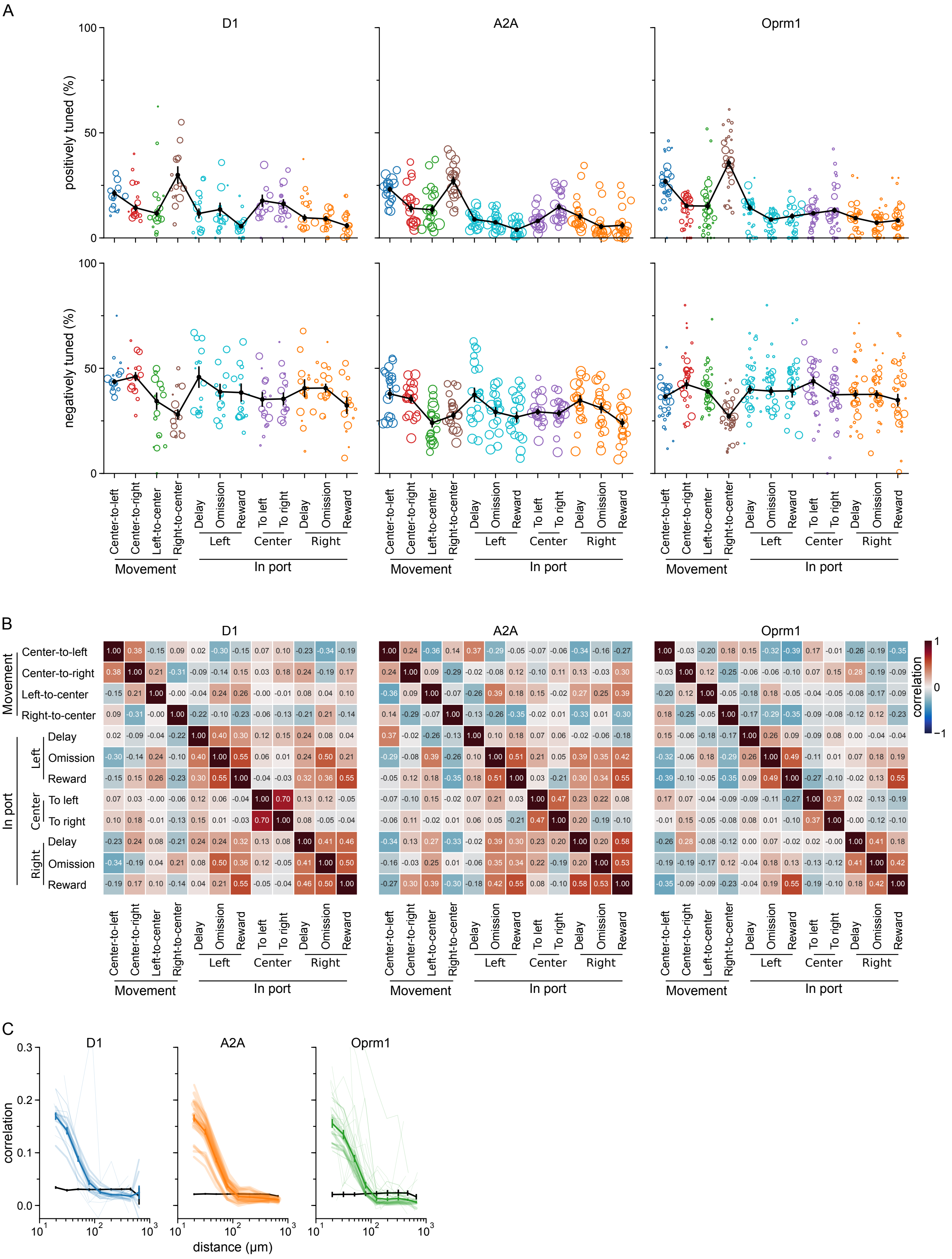
A
D1
A2A
Oprm1
positively tuned (%)
negatively tuned (%)
100
50
0
Center-to-left
Center-to-right
Left-to-center
Right-to-center
Delay
Omission
Reward
To left
To right
Left
Center
Right
Movement
In port
B
correlation
1
0
−1
C
correlation
0.3
0.2
0.1
0.0
distance (μm)

**Figure S4: Supplementary to Figure 3.** A) Fraction of neurons positively (increased firing) and negatively (decreased firing) tuned to the trial phases for each Cre-line. B) Correlations matrices of the tuning scores taken across all neurons of the respective Cre-line. C) The correlation between the binned (200ms) deconvolved signal of each pair of neurons versus the estimated distance between them. Each transparent line corresponds to one session and the opaque lines indicate averages weighted by the number of neurons in each session. Based on the first 30 minutes of each session.

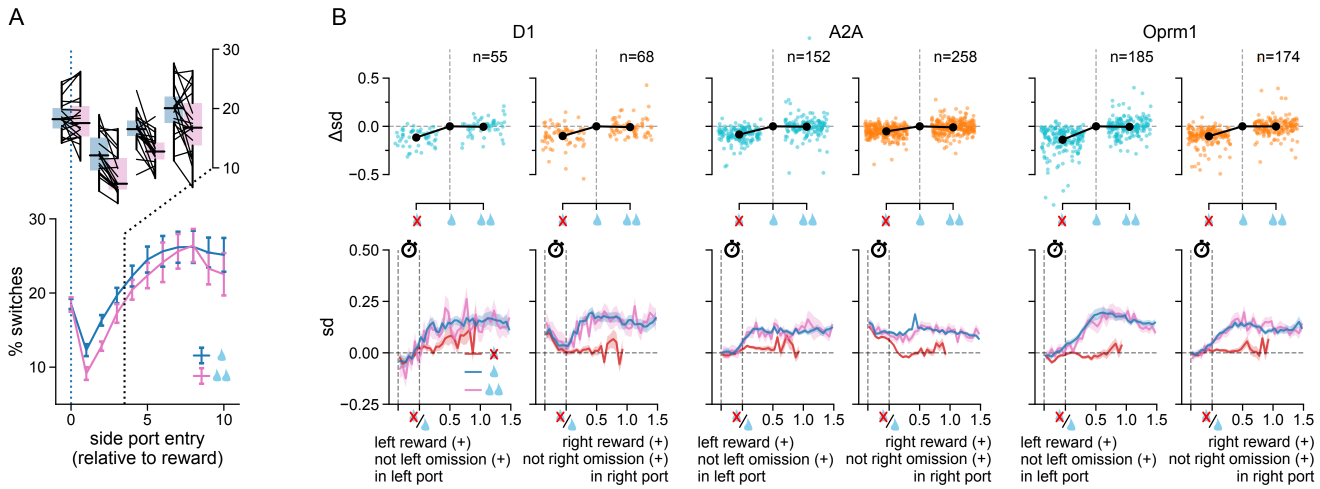


**Figure S5: Supplementary to Figure 3.** A) Doubling the size of the reward affects task performance. Mice are less likely to perform a port switch following the delivery of a doubled reward (pink, two drops) than following a single reward (blue, one drop). B) The activity of "reward neurons" (significantly tuned to the reward, but not, or negatively, tuned to the omission phase in choice ports) is not modulated by reward doubling. Top: Average response of left reward (cyan) and right reward (orange) neurons to reward omission (crossed-out drop) and reward doubling (two drops), relative to their average response to a single reward (one drop), in the left and right ports, respectively. Bottom: Frame-by-frame average deconvolved signal trace of the same neurons to in response to the same events.

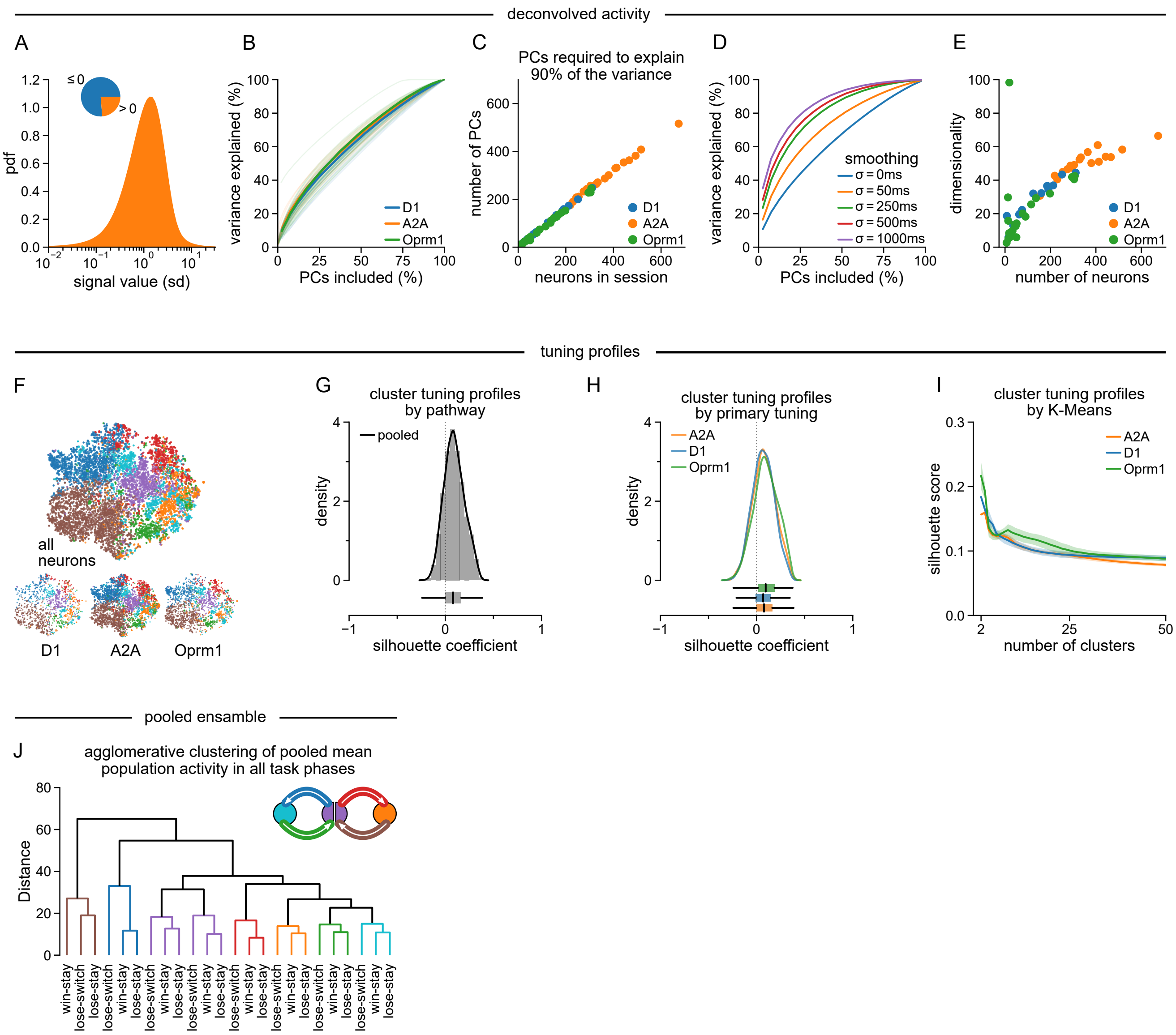


**Figure S6: Supplementary to Figure 3.** A) Histogram of std-normalized activity pooled over all Cre-lines, animals, sessions and neurons. Inset pie chart: the activity is mostly below 0 meaning no estimated spikes in the 50ms calcium imaging frame. Histogram: the value of the above-zero output form OASIS is heavy-tailed. B) A naive principal component analysis applied directly on the deconvolved traces from each session. The x-axis ranges from no principal components (PCs) to all PCs, i.e. the number of neurons. To compare across sessions with different amount of recorded neurons the number of PCs is normalized to the number of neurons. The y-axis indicate the fraction of variance that can be explained by the respective number of PCs (cumulative). Each thin line is one session and the thick lines are Cre-line averages. C) Same data as in B, but instead plotted as the number of PCs required to explain 90% of the variance. This seems to be close to 90% of the total number of PCs, suggesting the data cannot be projected to a lower-dimensional space without loss. D) By smoothing the deconvolved traces by a gaussian with increasing bandwidth the signal becomes a bit more low dimensional. However, the behaviors we are investigating are occurring at the 100ms timescale. E) Internal (non-linear) dimensionality also does not reveal any low dimensional manifolds. F-I) This analysis is performed on the tuning profiles, not the activity itself. F) tSNE of the tuning profiles for all neurons. Each neuron is represented by one point with the color of its primary tuning. G) The silhouette coefficient of each neuron assuming the cluster is given by the Cre-line. H) The silhoutte coefficient assuming the cluster identity is given by the primary tuning of each neuron. I) The average silhouette score assuming the cluster is given by a K-means clustering of the tuning profiles. J) Agglomerative clustering of pooled means. In particular, we calculated the average activity for each trial phase in each trial type and created a pooled "superensamble" with all neurons from all sessions. We then clustered the average activity vectors of the $8 \cdot 3 = 24$ phases assuming an euclidean distance metric. The trial phase (color) was always the first level of the split, followed by the trial type (x-label).

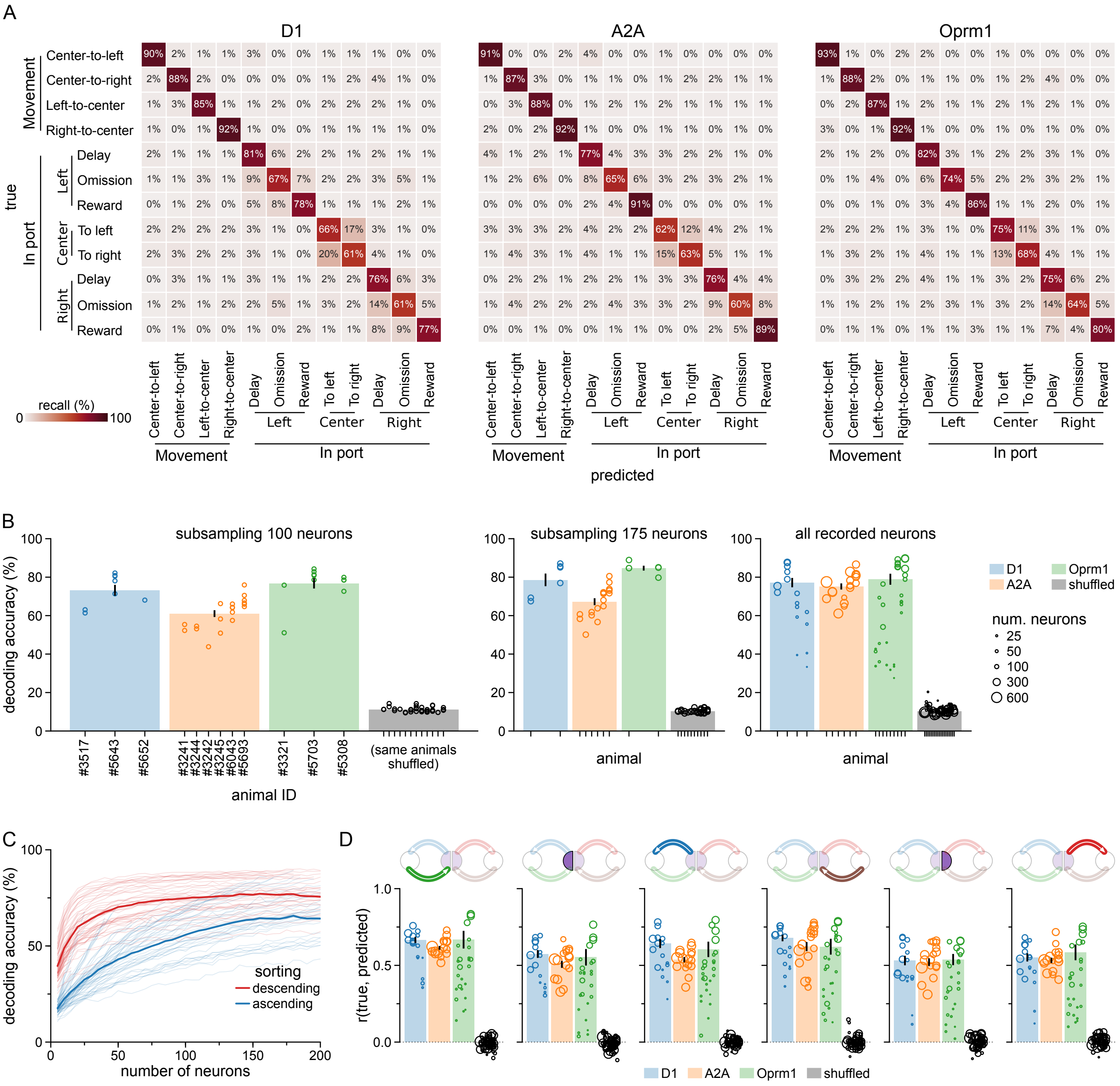


**Figure S7: Supplementary to Figure 3.** A) Full confusion matrix of the decoding of the trial phases, weighted by the number of neurons in each session. B) Data from Figure 3J split per animal at 100 neurons, 175 neurons and without subsampling. Each circle shows the cross-validated decoding accuracy during one recording session. Error bars show SEM calculated with bootstrapping. The means underlying the bars in the rightmost panel are weighted by the number of neurons in each session (size of the circle). The median decoding accuracy per animal was not significantly different between the pathways (Kruskal-Wallis H-test; at 100 neurons: N=3,6,3, H=5.14, p=0.076; at 175 neurons: N=2,6,2, H=4.98, p=0.082; at *all neurons*: N=4,6,9, H=2.44, p=0.29). C) Similar to Figure 3J, but rather than sampling neurons randomly they were sorted according to their mutual information with the trial phase (see Methods). Descending sorting imply choosing the "best" neurons first, ascending sorting imply choosing the "worst" neurons first. All sessions from all Cre-lines were included. D) The correlation between true and predicted progress through the trial phase (cf. Figure 3N) for six different trial phases. X-coordinate and circle sizes are as in B. Means are weighted by the number of neurons in each session.

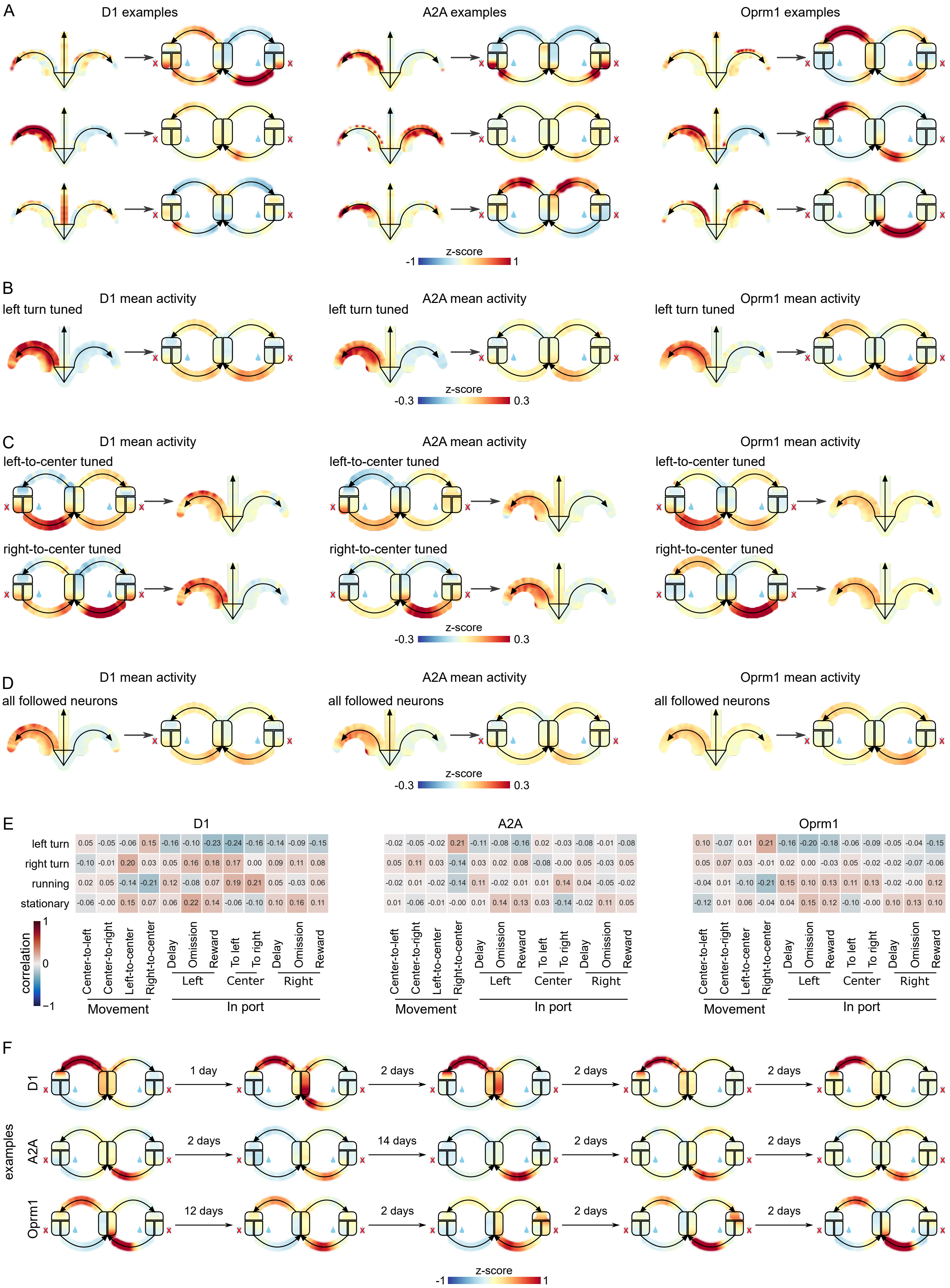
A
D1 examples
A2A examples
Oprm1 examples
z-score
-1
1
B
left turn tuned
D1 mean activity
A2A mean activity
Oprm1 mean activity
z-score
-0.3
0.3
C
left-to-center tuned
right-to-center tuned
D1 mean activity
A2A mean activity
Oprm1 mean activity
z-score
-0.3
0.3
D
all followed neurons
D1 mean activity
A2A mean activity
Oprm1 mean activity
z-score
-0.3
0.3
E
D1
left turn 0.05 -0.05 -0.06 0.15 -0.16 -0.10 -0.23 -0.24 -0.16 -0.14 -0.09 -0.15
right turn -0.10 -0.01 0.20 0.03 0.05 0.16 0.18 0.17 0.00 0.09 0.11 0.08
running 0.02 0.05 -0.14 -0.21 0.12 -0.08 0.07 0.19 0.21 0.05 -0.03 0.06
stationary -0.06 -0.00 0.15 0.07 0.06 0.22 0.14 -0.06 -0.10 0.10 0.16 0.11
A2A
-0.02 -0.05 -0.02 0.21 -0.11 -0.08 -0.16 0.02 -0.03 -0.08 -0.01 -0.08
0.05 0.11 0.03 -0.14 0.03 0.02 0.08 -0.08 -0.00 0.05 -0.03 0.05
-0.02 0.01 -0.02 -0.14 0.11 -0.02 0.01 0.01 0.14 0.04 -0.05 -0.02
0.01 -0.06 -0.01 -0.00 0.01 0.14 0.13 0.03 -0.14 -0.02 0.11 0.05
Oprm1
0.10 -0.07 0.01 0.21 -0.16 -0.20 -0.18 -0.06 -0.09 -0.05 -0.04 -0.15
0.05 0.07 0.03 -0.01 0.02 0.00 -0.03 0.05 -0.02 -0.02 -0.07 -0.06
-0.04 0.01 -0.10 -0.21 0.15 0.10 0.13 0.11 0.13 -0.02 -0.00 0.12
-0.12 0.01 0.06 -0.04 0.04 0.15 0.12 -0.10 -0.00 0.10 0.13 0.10
correlation
1
0
−1
Center-to-left
Center-to-right
Left-to-center
Right-to-center
Movement
Delay
Omission
Reward
Left
To left
To right
Center
Delay
Omission
Reward
Right
In port
F
examples
D1
1 day
2 days
2 days
2 days
A2A
2 days
14 days
2 days
2 days
Oprm1
12 days
2 days
2 days
2 days
z-score
-1
1

**Figure S8: Supplementary to Figure 4.** A) Nine more examples of neurons followed from the open field to the 2-choice task. B) The mean activity of all neurons tuned to left turns in the open field. C) The mean activity of all neurons tuned to the return movements in the 2-choice. D) For reference, the mean of all neurons that were followed from the open field to the 2-choice task. Note that the scale is different from individual examples. E) The correlations between the tuning scores in the open field and in the 2-choice task. F) Three example neurons aligned over five 2-choice sessions.

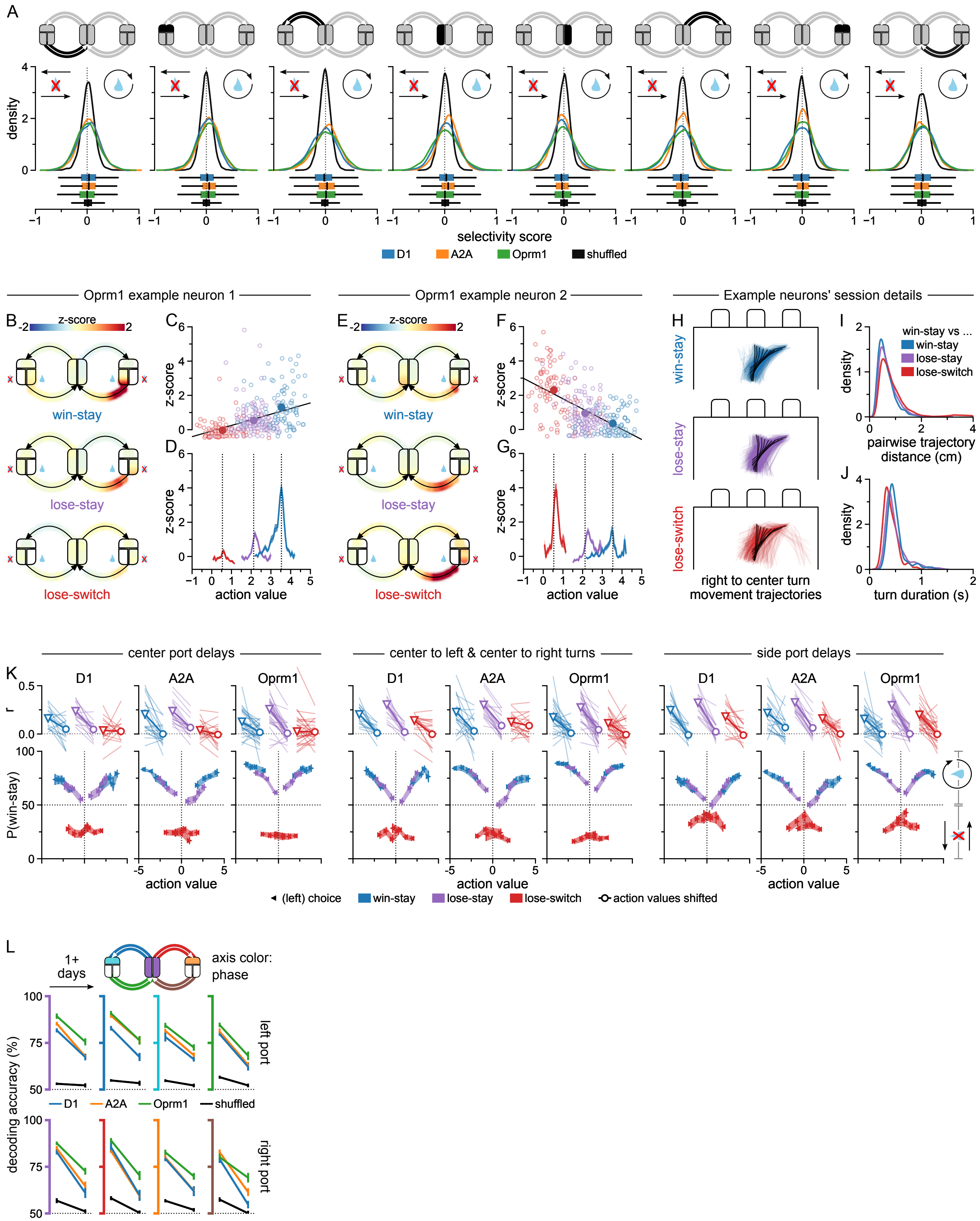
A
density
selectivity score
D1
A2A
Oprm1
shuffled
Oprm1 example neuron 1
Oprm1 example neuron 2
Example neurons' session details
B
z-score
win-stay
lose-stay
lose-switch
C
D
z-score
action value
E
F
G
H
right to center turn movement trajectories
I
win-stay vs ...
pairwise trajectory distance (cm)
J
turn duration (s)
center port delays
center to left & center to right turns
side port delays
K
r
P(win-stay)
(left) choice
action values shifted
L
1+ days
axis color: phase
decoding accuracy (%)
left port
right port

**Figure S9: Supplementary to Figure 5.** A) Distributions of neurons' phase-specific win-stay versus lose-switch selectivity scores, by Cre-line, for all trial phases except the outcome phase. A positive score indicates a neuron's preferential activation on win-stay, a negative score on lose-switch trials. Selectivity scores: area under the ROC curve, scaled from -1 to 1. For comparison, selectivity scores were calculated using shuffled behavior data (black). n=1943 D1+ neurons, 6566 A2A+ neurons, 2793 Oprm1+ neurons. B) Example Oprm1+ neuron highly selective for win-stay over lose-switch trials in the right port to center turn phase. The intermediate response on lose-stay trials suggests the selectivity may be a consequence of action value coding. C) The average activity of the neuron in B during right port to center turns correlates with action value. Open circles: single trials. Filled circles: average of single trials by trial type. Color-coding indicates trial type as in B. Black line: least-squares regression line. D) Average activity over time (2 sec window) of the same neuron as in B and C, aligned to right port-exits (dotted line), by trial type. Despite the x-axis unit (action value), the traces represent a time series. On the x-axis, the traces are centered on the mean action value of the averaged trials. Color-coding as in B and C. Shading: ±SEM. E-G) Same as B-D, but for a lose-switch selective Oprm1+ neuron recorded in the same session. H) Frame-by-frame right port to center turn movement trajectories by trial type, recorded in the same session as the example neurons in B-G. Average trajectories in black (10% wide turn progress bins). The mouse is represented, frame-for-frame, by a line connecting the base of the tail, the center of the body, and the point between the ears. I) Density plot of the pairwise distances (in cm) between the single trial-turn trajectories shown in H. We compared individual win-stay trajectories to lose-stay, lose-switch, and all other win-stay trajectories and found the distances to be similarly distributed for all three comparisons. To compare single trials, we first binned each trial by turn progress (10% wide bins) and computed the average position of each tracked point per bin in order to standardize trial length. For each pair of trials, we then computed the average distance of the 3 tracked points (tail base, body, head) bin-for-bin, and finally obtained the average over all 10 bins as the trajectory distance. J) Density plot of the turn durations (in sec) of the turns shown in H, by trial type. K) Correlations of SVMs' trial-by-trial win-stay (versus lose-switch) probability estimates with trial action value for the center port, center-to-side-port turn, and side port delay trial phases, by trial type (i.e. win-stay, lose-stay and lose-switch) and Cre-line. Top panels: actual probability–action value correlations compared to correlations computed using randomly shifted action values. Individual sessions (thin lines) and average of sessions weighted by number of neurons (thick lines) shown. Bottom panels: win-stay probability estimates as a function of action value. Triangles: mean probability estimate of sessions weighted by number of neurons. Shading and error bars: ±SEM (bootstrapped). L) Decoding accuracy of win-stay versus lose-switch trial type-predicting SVMs for training-session data compared to their accuracy when applied to test-session data recorded one or more days later, by trial phase (axes color-coded) and Cre-line (line color). Black lines: average accuracy of decoders trained on shuffled data. Averages of pairs of sessions weighted by the number of neurons matched across sessions. Error bars: ±SEM (bootstrapped).

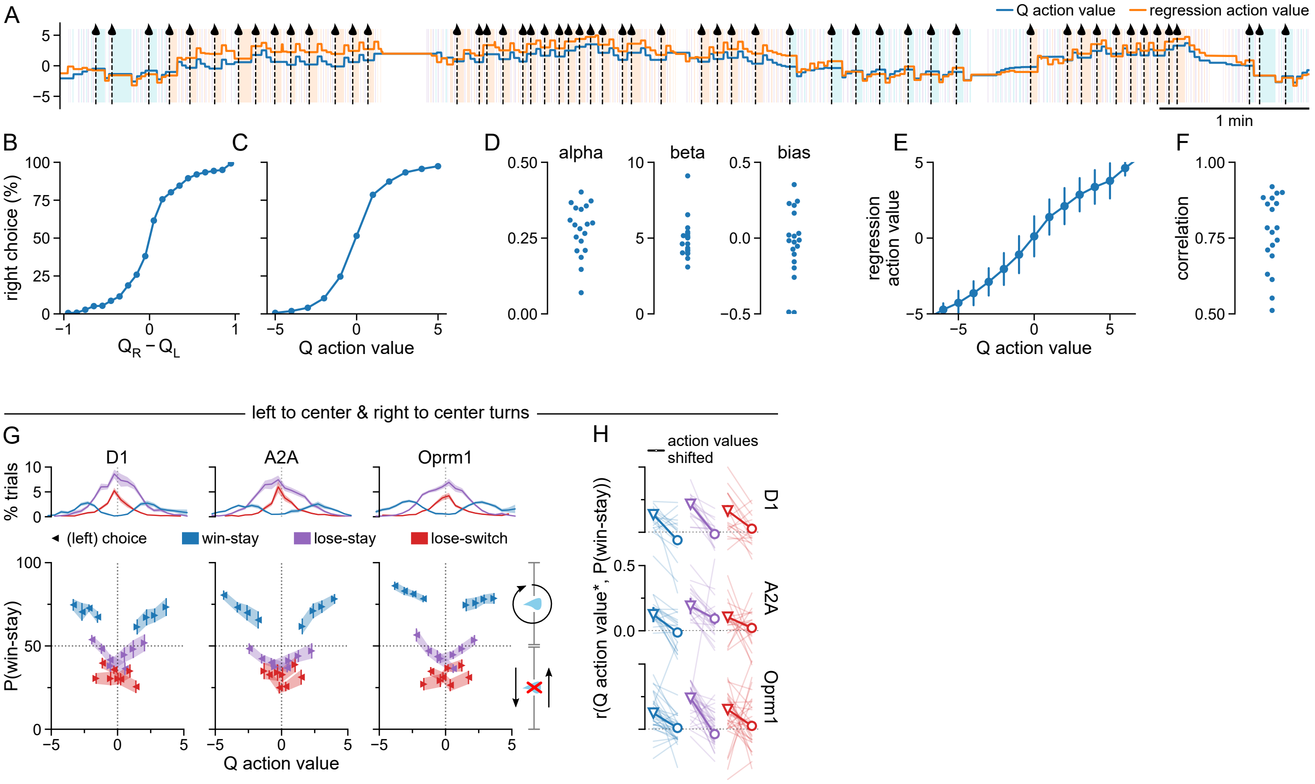


**Figure S10: Supplementary to Figure 5.** A) Example comparison of the two ways of calculating action values. Background colors and symbols follow Figure 3A. B) Probability of choosing the right port as a function of the difference between the Q-values. Data pooled from all animals. C) Probability of choosing the right port as a function of Q action value (i.e. $\beta(Q_R - Q_L) + \text{bias}$). D) The three fitted parameters (see Methods) for each animal. E) Comparison between the two ways of calculating action values. Error bars indicate standard deviation. F) Correlation between Q action value and regression action value for each animal. G) Replication of Figure 5P but with Q-action value. H) Replication of Figure 5Q but with Q-action value.

| animal ID | date | task | duration | neurons | trials |
|---|---|---|---|---|---|
| 3517 | 2018-03-29 | 2-choice | 60m17s | 311 | 637 |
| | 2018-04-04 | 2-choice | 64m17s | 212 | 918 |
| 5643 | 2019-01-12 | 2-choice | 60m14s | 251 | 732 |
| | 2019-01-14 | 2-choice | 60m16s | 189 | 866 |
| | 2019-01-28 | 2-choice | 61m22s | 185 | 838 |
| | 2019-01-30 | Open field | 20m1s | 191 | |
| | | 2-choice | 21m2s | 191 | 243 |
| | 2019-02-01 | Open field | 15m6s | 160 | |
| | | 2-choice | 45m1s | 160 | 636 |
| | 2019-02-24 | Open field | 24m23s | 156 | |
| 5651 | 2019-01-31 | 2-choice | 61m44s | 62 | 506 |
| | 2019-02-03 | 2-choice | 60m51s | 36 | 954 |
| | 2019-02-04 | 2-choice | 62m27s | 19 | 884 |
| | 2019-02-05 | 2-choice | 50m26s | 8 | 954 |
| | 2019-02-24 | Open field | 21m5s | 24 | |
| 5652 | 2019-01-28 | 2-choice | 62m33s | 124 | 666 |
| | 2019-01-30 | 2-choice | 61m34s | 75 | 639 |
| | 2019-01-31 | 2-choice | 60m59s | 65 | 569 |
| | 2019-02-02 | Open field | 15m10s | 40 | |
| | | 2-choice | 45m1s | 40 | 602 |
| | 2019-02-03 | Open field | 15m22s | 15 | |
| | | 2-choice | 45m23s | 15 | 577 |
| | 2019-02-24 | Open field | 22m45s | 46 | |

**Table S1:** All *recorded* sessions with D1 animals.

| animal ID | date | task | duration | neurons | trials |
|---|---|---|---|---|---|
| 3241 | 2018-03-26 | 2-choice | 60m6s | 674 | 689 |
| | 2018-04-03 | 2-choice | 63m0s | 380 | 536 |
| 3242 | 2018-03-30 | 2-choice | 60m5s | 466 | 315 |
| 3244 | 2018-03-30 | 2-choice | 61m30s | 443 | 598 |
| | 2018-04-05 | 2-choice | 61m52s | 413 | 453 |
| 3245 | 2018-04-03 | 2-choice | 60m3s | 364 | 533 |
| | 2018-04-05 | 2-choice | 60m55s | 154 | 598 |
| | 2018-04-10 | 2-choice | 60m8s | 219 | 567 |
| 5693 | 2019-01-15 | 2-choice | 61m43s | 408 | 912 |
| | 2019-01-16 | 2-choice | 64m55s | 334 | 997 |
| | 2019-01-27 | 2-choice | 60m1s | 295 | 596 |
| | 2019-01-29 | 2-choice | 66m5s | 254 | 606 |
| | 2019-01-31 | Open field | 14m21s | 275 | |
| | | 2-choice | 45m13s | 275 | 789 |
| | 2019-02-02 | Open field | 16m5s | 304 | |
| | | 2-choice | 40m5s | 304 | 877 |
| | 2019-02-24 | Open field | 20m46s | 277 | |
| 6043 | 2019-01-14 | 2-choice | 60m6s | 328 | 895 |
| | 2019-01-26 | 2-choice | 57m23s | 493 | 798 |
| | 2019-01-28 | 2-choice | 59m14s | 230 | 1064 |
| | 2019-01-30 | Open field | 15m2s | 288 | |
| | | 2-choice | 45m2s | 288 | 912 |
| | 2019-02-01 | Open field | 15m11s | 244 | |
| | | 2-choice | 46m15s | 244 | 1102 |
| | 2019-02-24 | Open field | 20m14s | 216 | |

**Table S2:** All *recorded* sessions with A2A animals.

| animal ID | date | task | duration | neurons | trials |
|---|---|---|---|---|---|
| 3321 | 2018-03-27 | 2-choice | 60m39s | 136 | 539 |
| | 2018-03-31 | 2-choice | 60m34s | 78 | 414 |
| | 2018-04-09 | 2-choice | 61m18s | 115 | 984 |
| 3323 | 2018-03-27 | 2-choice | 60m4s | 33 | 494 |
| | 2018-03-31 | 2-choice | 60m27s | 29 | 426 |
| | 2018-04-09 | 2-choice | 60m1s | 47 | 815 |
| 3572 | 2018-03-29 | 2-choice | 60m11s | 18 | 464 |
| | 2018-04-03 | 2-choice | 60m24s | 19 | 670 |
| 3581 | 2018-04-02 | 2-choice | 60m47s | 13 | 391 |
| 3582 | 2018-03-27 | 2-choice | 60m52s | 17 | 320 |
| | 2018-03-29 | 2-choice | 60m12s | 50 | 567 |
| | 2018-04-04 | 2-choice | 60m24s | 20 | 340 |
| 5308 | 2019-01-31 | 2-choice | 67m42s | 304 | 1152 |
| | 2019-02-01 | Open field | 15m3s | 295 | |
| | | 2-choice | 45m2s | 295 | 783 |
| | 2019-02-04 | Open field | 15m2s | 307 | |
| | | 2-choice | 42m39s | 307 | 633 |
| | 2019-02-05 | 2-choice | 50m53s | 138 | 933 |
| | 2019-02-06 | 2-choice | 40m55s | 90 | 703 |
| | 2019-02-24 | Open field | 20m6s | 236 | |
| 5464 | 2019-01-14 | 2-choice | 60m3s | 15 | 963 |
| | 2019-02-05 | 2-choice | 60m34s | 14 | 887 |
| | 2019-02-07 | 2-choice | 59m23s | 7 | 586 |
| | 2019-02-24 | Open field | 20m4s | 12 | |
| 5574 | 2019-01-26 | 2-choice | 63m16s | 43 | 897 |
| | 2019-01-27 | 2-choice | 60m20s | 51 | 855 |
| | 2019-01-29 | 2-choice | 60m16s | 42 | 823 |
| | 2019-01-31 | Open field | 15m40s | 27 | |
| | | 2-choice | 45m4s | 27 | 759 |
| | 2019-02-02 | Open field | 15m21s | 26 | |
| | | 2-choice | 45m32s | 26 | 736 |
| | 2019-02-24 | Open field | 22m16s | 62 | |
| 5703 | 2019-01-14 | 2-choice | 60m11s | 198 | 801 |
| | 2019-01-16 | 2-choice | 61m6s | 162 | 873 |
| | 2019-01-26 | 2-choice | 60m14s | 169 | 755 |
| | 2019-01-30 | 2-choice | 45m16s | 181 | 800 |
| | 2019-01-30 | Open field | 15m1s | 181 | |
| | 2019-02-01 | 2-choice | 45m2s | 149 | 700 |
| | 2019-02-01 | Open field | 15m2s | 149 | |
| | 2019-02-24 | Open field | 29m21s | 170 | |

**Table S3:** All *recorded* sessions with Oprm1 animals.
